# Widening the landscape of transcriptional regulation of green algal photoprotection

**DOI:** 10.1101/2022.02.25.482034

**Authors:** Marius Arend, Yizhong Yuan, M. Águila Ruiz-Sola, Nooshin Omranian, Zoran Nikoloski, Dimitris Petroutsos

**Affiliations:** Bioinformatics Group, Institute of Biochemistry and Biology, University of Potsdam, 14476 Potsdam, Germany; Systems Biology and Mathematical Modeling Group, Max-Planck-Institute of Molecular Plant Physiology, 14476 Potsdam, Germany; Bioinformatics and Mathematical Modeling Department, Center of Plant Systems Biology and Biotechnology, 4000 Plovdiv, Bulgaria; University of Grenoble Alpes, CNRS, CEA, INRAE, IRIG-LPCV, 38000 Grenoble, France

## Abstract

Availability of light and CO2, substrates of microalgae photosynthesis, is frequently far from optimal. Microalgae activate photoprotection under strong light, to prevent oxidative damage, and the CO2 Concentrating Mechanism (CCM) under low CO2, to raise intracellular CO2 levels. The two processes are interconnected; yet, the underlying transcriptional regulators remain largely unknown. Employing a large transcriptomics data compendium of *Chlamydomonas reinhardtii’s* responses to different light and carbon supply, we reconstructed a consensus genome-scale gene regulatory network from complementary inference approaches and used it to elucidate transcriptional regulators of photoprotection. We showed that the CCM regulator LCR1 also controls photoprotection, and that QER7, a Squamosa Binding Protein, suppresses photoprotection- and CCM-gene expression under the control of the blue light photoreceptor Phototropin. By demonstrating the existence of regulatory hubs that channel light- and CO2-mediated signals into a common response, our study provides an accessible resource to dissect gene expression regulation in this microalga.

## Introduction

Photosynthetic microalgae convert light into chemical energy in the form of ATP and NADPH to fuel the CO_2_ fixation in the Calvin–Benson cycle^1^. They have evolved to cope with rapid fluctuations in light^2^ and inorganic carbon (Ci)^3^ availability in their native habitats. When absorbed light exceeds the CO_2_ assimilation capacity, the formation of harmful reactive oxygen species can lead to severe cell damage; this is prevented by the activation of photoprotective mechanisms, collectively called non-photochemical quenching (NPQ). NPQ encompasses several processes that are distinguished in terms of their timescales^2^, among which the rapidly reversible energy-quenching (qE) is, under most circumstances, the predominant NPQ component^2, 4^. The major molecular effector of qE in the green model microalga *Chlamydomonas reinhardtii* (hereafter *Chlamydomonas*) is the LIGHT HARVESTING COMPLEX STRESS RELATED protein LHCSR3, encoded by the *LHCSR3.1* and *LHCSR3.2* genes^5^ that slightly differ only in their promoters; LHCSR1 can also contribute significantly to qE under conditions where LHCSR3 is not expressed^6, 7^. PSBS, the key qE effector protein in higher plants^8^ is encoded in two highly similar paralogues *PSBS1* and *PSBS2* in *Chlamydomonas*^9^. They are only transiently expressed in *Chlamydomonas* under high light (HL)^9, 10^ and their gene products accumulate under UV-B irradiation^11^; while their precise contribution in *Chlamydomonas* photoprotective responses is still a matter of ongoing research, current understanding is that PSBS proteins contribute to photoprotection during HL acclimation of *Chlamydomonas* through both NPQ-independent and NPQ-dependent mechanisms^12^.

Intracellular levels of CO_2_ are modulated by the availability of its gaseous and hydrated forms^3^ in the culture media and the supply of acetate, that is partly metabolized into CO2^7, 13^. Under low CO_2_, *Chlamydomonas* activates the CO_2_-concentrating mechanism (CCM) to avoid substrate-limitation of photosynthesis by raising the CO_2_ concentration at the site of RuBisCO, where CO_2_ is assimilated^3^. The CCM mainly comprises of carbonic anhydrases (CAHs) and of inorganic carbon transporters. Almost all CCM-related genes are under the control of the nucleus-localized zinc-finger type nuclear factor CIA5 (aka CCM1)^14–16^, including the Myb Transcription Factor LOW-CO_2_ -STRESS RESPONSE 1 (LCR1) that controls expression of genes coding for the periplasmic CAH1, the plasma membrane-localized bicarbonate transporter LOW CO_2_-INDUCED 1 (LCI1), and the low-CO_2_ responsive LCI6, whose role remains to be elucidated^17^. CIA5 is also a major qE regulator activating transcription of genes encoding LHCSR3 and PSBS, while repressing accumulation of LHCSR1 protein^7^.

LHCSR3 expression relies on blue light perception by the photoreceptor phototropin (PHOT)^18^, on calcium signaling, mediated by the calcium sensor CAS^19^ and on active photosynthetic electron flow^18–20^, likely via indirectly impacting CO_2_ availability^7^. The critical importance of CO2 in LHCSR3 expression is demonstrated by the fact that changes in CO2 concentration can trigger LHCSR3 expression^21–23^ even in the absence of light^7^. Accumulation of *LHCSR1* and *PSBS* mRNA is under control of the UV-B photoreceptor UVR8^11^ and PHOT^24, 25^ and is photosynthesis-independent^20, 25^. While LHCSR1 is CO2/CIA5 independent at the transcript level^7, 25^, PSBS is responsive to CO2 abundance and is under partial control of CIA5^7^. A Cullin (CUL4) dependent E3-ligase^24, 26, 27^ has been demonstrated to post-translationally regulate the transcription factor (TF) complex of CONSTANS (CrCO)^26^ and NF-Y isomers^27^, which bind to DNA to regulate the transcription of *LHCSR1*, *LHCSR3*, and *PSBS*. The putative TF and diurnal timekeeper RHYTHM OF CHLOROPLAST 75 (ROC75) was shown to repress LHCSR3 under illumination with red light^28^.

Here, we employed a large compendium of RNAseq data from *Chlamydomonas* to build a gene regulatory network (GRN) underlying light and carbon responses, and thus reveal the transcriptional regulation of qE at the interface of these responses. The successful usage of RNAseq data to infer GRNs has been demonstrated in many studies^29–31^, although the data pose some challenges that require careful consideration. All of the developed approaches to infer a GRN quantify the interdependence between the transcript levels of TF-coding genes and their putative targets; the resulting prediction model serves as a proxy for the regulatory strength that the product of the TF-coding gene exerts on its target(s). It is usually the case that the number of observations (samples) used in building the model is considerably smaller than the number of TFs used as predictors, leading to collinearity of the transcript levels and associated computational instabilities; furthermore, as an artifact of the computational techniques, some of the inferred regulations may be spurious^32–34^. To address these issues, here we took advantage of combining the outcome of multiple regularization techniques and post-processing to increase the robustness of identified interactions^33–35^. In contrast to our approach, the existing predicted GRNs of *Chlamydomonas* either focused on nitrogen starvation^36^ or used a broad RNAseq data compendium, not tailored to inferring regulatory interactions underlying responses to particular cues^37^. Moreover, these GRNs were not obtained by combining the outcomes from multiple inference approaches, shown to increase accuracy of predictions^30^, and their quality was not gauged against existing knowledge of gene regulatory interactions.

We used an RNAseq data compendium of 158 samples (**Supplementary Table 1**) from *Chlamydomonas* cultures exposed to different light and carbon supply as input to seven benchmarked GRN inference approaches that employ complementary inference strategies^29, 30^ to identify activating and inhibiting regulatory interactions. We assessed the performance of each approach based on a set of curated TF-target gene interactions with experimental evidence from *Chlamydomonas.* Based on this assessment, we integrated the outcome of the five best performing approaches into a unique resource, a consensus network of *Chlamydomonas* light- and carbon-dependent transcriptional regulation. We used the consensus network to reveal regulators of qE genes and demonstrated the quality of predictions by experimentally validating two of the six tested candidates. We show here that LCR1 regulates not only CCM, as previously reported^17^, but also qE by activating the expression of LHCSR3, and demonstrate that qE-REGULATOR 7 (QER7), belonging to the SQUAMOSA-PROMOTER BINDING PROTEIN-LIKE gene family, is a repressor of qE and CCM-gene expression. Our work consolidates the extensive co-regulation of CCM and photoprotection^7^ based on the untargeted assessment of the obtained genome-scale GRN.

## Results

### Computationally inferred GRN recovered known regulatory interactions underlying qE and CCM in Chlamydomonas

We first aimed to employ published and in-house generated RNAseq data sets capturing the transcriptional responses of *Chlamydomonas* to light and acetate availability to infer the underlying GRN. To this end, we obtained data from two publicly available transcriptomics studies of synchronized chemostat wild-type (WT) cultures grown in a 12h/12h light dark scheme and sampled in 30 min to 2h intervals^38, 39^. We combined these with our RNAseq data generated from mixotrophically or autotrophically grown batch cultures of the WT and *phot* mutant acclimated to low light (LL) or exposed to HL (**Methods, Supplementary Table 1**). These data sets capture the expected expression patterns of the key genes involved on CCM and qE (**Fig. 1a, Supplementary Fig. 1**) in response to changes in acetate availability and light intensity. Specifically, we found strong up-regulation of these genes in the light^7, 25^, and a marked inhibition of *LHCSR3.1/2* and CCM genes by acetate as previously described^7, 40^.

**Fig. 1:**
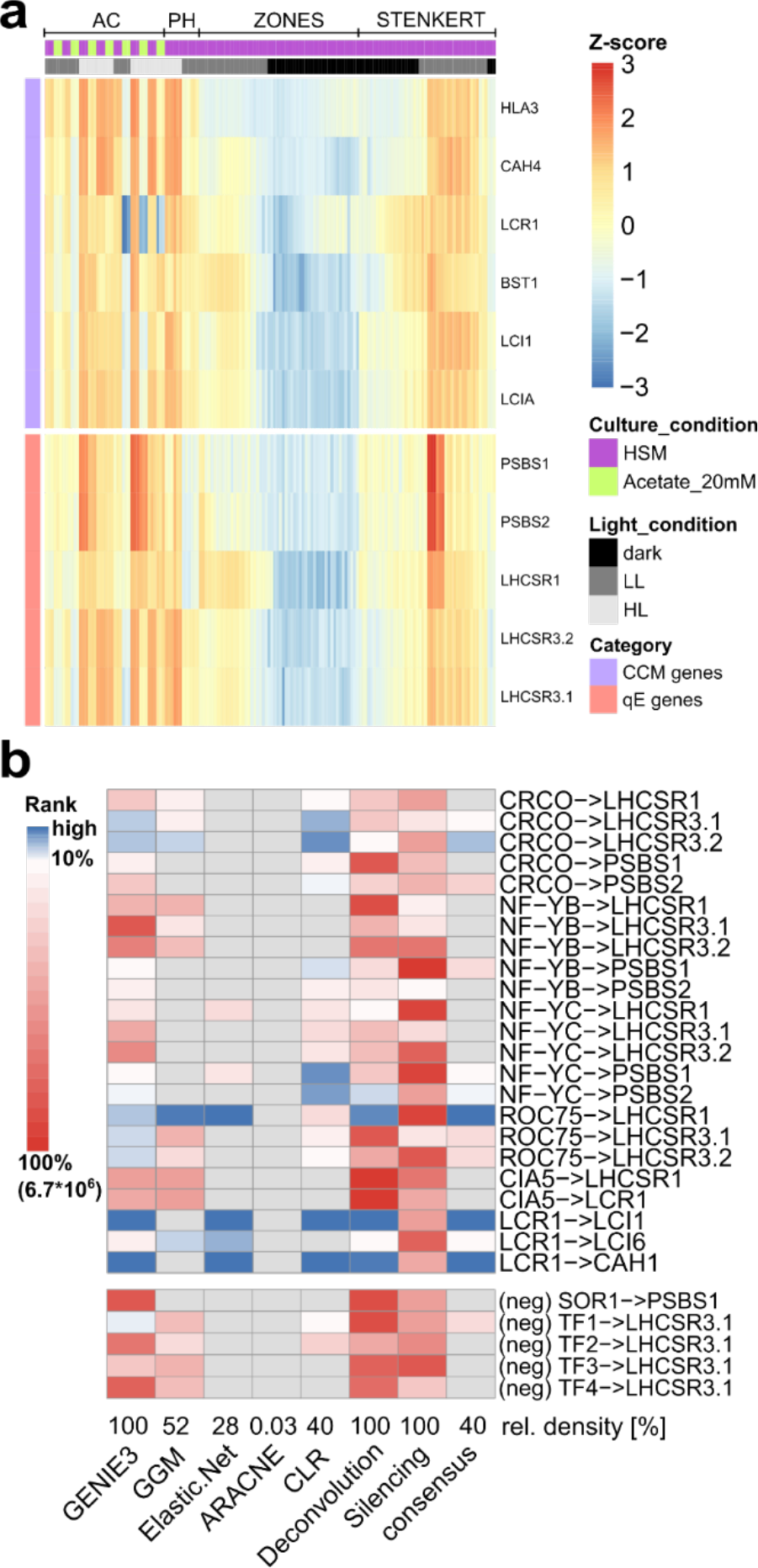
Characterization of the consensus GRN inferred by employing a compendium of RNA-seq data from diverse light and culture conditions. A. Expression levels of representative CCM and qE genes are plotted over all samples used for network inference (z-scaled log values are depicted). The column annotation gives information on the culture conditions and data set (see also Supplementary Table 1); no clustering was applied. Supplementary Fig. 1 shows an enlarged version of this figure with sample names included. B. The heatmap rows correspond to experimentally validated or invalidated (neg) gene regulatory interactions involved in qE and CCM, curated from literature. The heatmap indicates ranking of these interactions by different approaches and the consensus network. Edges are considered highly ranked (depicted in blue) if they are above the 10% network density threshold. Edges ranked below this threshold are depicted in red. Edges that were not included in the given network are marked in grey. At the bottom of the heatmap the relative number of edges inferred by each approach is provided. ARACNE and Silencing columns were only plotted for comparison and were not used in building the consensus GRN (see Methods). TF1: MYB-like DNA- binding protein, Cre01.g034350; TF2: RWP8, RWP-RK transcription factor, Cre04.g218050; TF3: RWP5, RWP- RK transcription factor, Cre06.g285600; TF4: bHLH domain-containing protein, Cre07.g349152)

We employed these data together with a list of 407 transcription factors from protein homology studies^41, 42^ **(Methods, Supplementary Table 2)**, as input to seven GRN inference approaches to robustly predict TF-target interactions, as shown in benchmark studies^30^. The employed methods infer both activating and inhibiting TF- target interactions. Benchmarking of GRNs usually relies on ground truth data obtained from ChIPseq or transcriptomic profiling of TF mutants. Since to date no such comprehensive data set exists for *Chlamydomonas* we manually curated a list of known, experimentally validated regulatory interactions underlying CCM and qE in *Chlamydomonas*^22, 26–28^, to assess the quality of GRNs inferred by the different approaches. As negative control, we included the reported lack of effect of the SINGLET OXYGEN RESISTANT 1 (SOR1) TF on *PSBS1* transcript levels in diurnal culture^43^ as well as a set of four TFs, designated TF1-4, whose knock-out or overexpression did not affect *LHCSR3.1* transcript levels (**Supplementary Note 1**, **Supplementary Fig. 2**, **Fig. 1b**). When we assessed the predicted ranks, as a measure of confidence assigned to the positive and negative ground truth data, we found that two approaches were clear outliers, showing sensitivity of 0%. More specifically, ARACNE^44^ and global silencing^45^ are unable to recover any positive literature interactions when using a network density threshold of 10% of all possible TF-TF and TF-target interactions (most prominently observable in the case of LCR1 interactions). Possible reasons for this finding are over-trimming or issues with the validity of the underlying assumptions, as seen in other case studies^46^. Since the presence of most gene regulatory interactions is dependent on environmental stimuli, it is considerably easier to experimentally validate the presence of TF-target interactions than to show that they do not take place^47^. Thus, in inferring a consensus GRN we considered the five approaches that were able to recover positive interactions, namely: Graphical Gaussian Models (GGM), Context Likelihood of Relatedness (CLR), Elastic Net regression, Gene Network Inference with Ensemble of Trees (GENIE3), and Network Deconvolution (**Methods**).

To obtain insights in the performance of these approaches, we next quantified the variability of ranks for the known TF-target gene interactions. We found that the average standard deviation of the ranks of the TF- target gene interactions within an approach is larger than the average standard deviation for the rank of a TF- target gene interaction across the five approaches (**Supplementary Fig. 3**, **Fig. 1b**). This observation suggested that the properties of a given TF-target gene interaction have a stronger influence on its assigned rank than the inference approach used. More specifically, we noted that the regulation of LHCSR genes by the two NF- Y paralogues and the induction of LCR1 by CIA5 are not recovered by any of the used approaches; this is in line with reports showing that CIA5 is constitutively expressed and regulated post-translationally^16^ - not reflected in the transcriptomics data. Further, NF-Y factors that rely on complex formation with CrCO to regulate their targets^27^, may also act via unresolved post-translational mechanisms. As mentioned in the introduction, some of the employed approaches use regularization techniques, mitigating effects of collinearity and low sample number, this comes at the cost of increasing the number of false negatives, another reason for the high number of unrecovered interactions previously reported in literature. Importantly, the regulatory interactions of the CCM effector genes *LCI1* and *CAH1* by *LCR1* are assigned very high ranks (top 1%) by the approaches considered in the consensus GRN (except for GGM); moreover, only for the interaction of TF1 and LHCSR3.1 we observe false positives, originating from spurious interactions. The interactions of the other four TFs (**Supplementary Note 1, Supplementary Fig. 2**) are correctly discarded by all approaches, indicating the robustness of the employed approaches (**Fig. 1b**). In addition, we observed that CLR and GENIE3 demonstrated the best performance with respect to the set of known interactions. For instance, they identified the regulation of *LHCSR3.1* by CrCO^26, 27^ and of *PSBS1* by NF-YB^27^ (**Fig. 1b**).

Generalization of this ranking beyond the known interactions underlying qE and CCM processes is challenging, due to the lack of genome-scale gold standard, and we therefore opted to combine the results of the five approaches, that showed comparable performance, in the consensus GRN **(Methods, Supplementary Table 3)** to increase robustness of the predictions. Our analyses of the overlap between the consensus and individual GRNs and the enrichment of TF-TF interactions demonstrated the robustness of the inferred interactions **(Supplementary Note 2, Supplementary Fig. 4)**.

### Consensus GRN pinpoints LCR1 as a regulator of qE-related genes

Using the consensus GRN, we inferred direct regulators of *LHCSR* and *PSBS* genes and ranked them according to the score resulting from the Borda method (**Methods**)^30, 48^. Mutants were available for four of the top ten of TFs with strongest cumulative regulatory effect on qE-related genes (**Fig. 2a, Supplementary Table 4):** Two knock-out mutants of previously uncharacterized genes were ordered from the CliP library^49^, which we termed *qE-regulators 4* and *6* (*qer4*, *qer6*; see **Supplementary Fig. 5** for the genotyping of these mutants). Additionally, we obtained an over-expressor line of the N-acetyltransferase LCI8^21^ and the knock-out strain of the known CCM regulator LCR1^17^. We tested for a regulatory effect by switching LL-acclimated mutant strains and their respective WT background to HL for 1h and quantified transcript levels of qE-related genes. Since both the paralogs of *PSBS* as well as of *LHCSR3* show correlation greater than 0.96 over all RNAseq samples used in this study and they additionally have very similar expression profiles quantified by RT-qPCR^50^, we only probed the transcripts of *LHCSR3.1* and *PSBS1* via qPCR in the validation assays. For *qer4*, *qer6*, and *lci8-oe* we did not observe an effect on the transcript levels of investigated genes after HL exposure (**Supplementary Fig. 6**). Thus, qer4 and qer6 are considered false positive predictions of the GRN, despite the fact that *qer4* accumulated 1.5 times more *LHCSR3.1* under LL than the WT. A review of the closest orthologs of LCI8 together with the experimental data indicate that it is likely involved in arginine synthesis^51^ and wrongly included as histone acetylase in the list of TFs. Interestingly, LCR1, the highest ranking among the tested regulators showed significantly decreased expression of LHCSR3 at both the gene (three times lower, **Fig. 2b**) and protein level (four times lower, **Fig. 2c, d**) compared to the WT; as a result, *lcr1* developed very low NPQ and qE **(Fig. 2e)**. Interestingly, the *lcr1* mutant over-accumulated LHCSR1 and PSBS both at the transcript and at the protein level (**Fig. 2b-d**); Complementation of *lcr1* with the knocked-out gene (strain *lcr1-C*) restored LHCSR3 gene and protein expression as well as the qE phenotype (**Fig. 2b-e**). Because pre-acclimation conditions impact qE gene expression^25^ we conducted independent experiments in which cells were acclimated to darkness before exposure to HL and we obtained very similar results. Our data demonstrated that *lcr1* showed significantly lower expression of LHCSR3 and higher expression of LHCSR1/PSBS at both the gene (**Supplementary Fig. 7a**) and protein level (**Supplementary Fig. 7b,c**), and had lower NPQ and qE (**Supplementary Fig. 7d**) than the WT, although the higher expression levels of the *LHCSR1* gene were not rescued by the complementation with the missing *LCR1* gene (**Supplementary Fig. 7a**). Altogether, our data show that LCR1 is a regulator of qE by activating *LHCSR3.1* transcription and repressing LHCSR1 and PSBS accumulation.

**Fig. 2:**
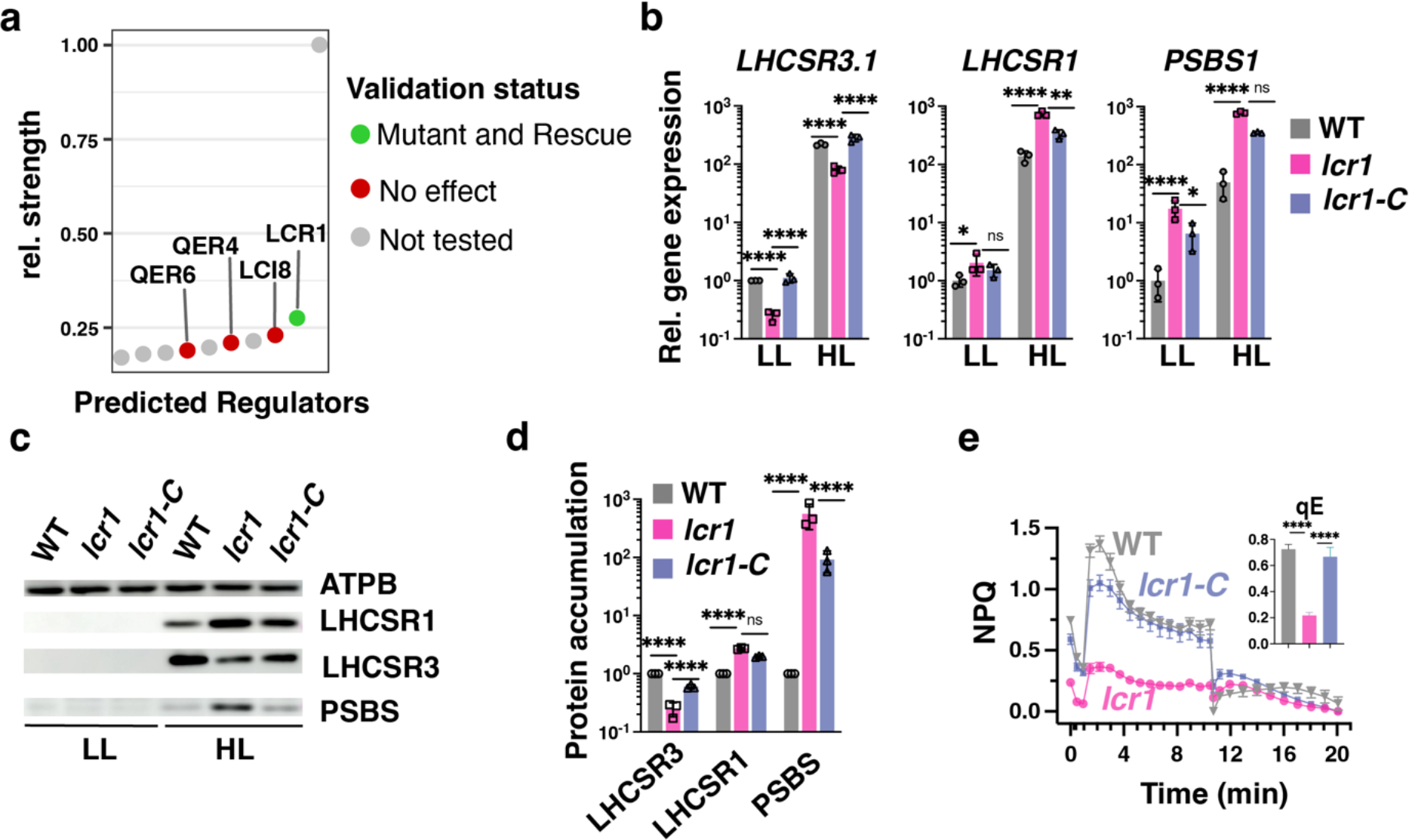
Consensus GRN for light and acetate responses pinpoints LCR1 as regulator of qE-related genes. **A.** Dot plot of the relative regulatory strength of the top 10 regulators of qE-related genes in the consensus GRN (see Methods). TFs are marked in green if qE transcript levels were affected in the respective knock-out strain and this effect was reversed by complementation with the missing gene. TFs for which no effect was observed are marked in red. TFs for which no mutant lines were available are plotted in grey. **B.** WT, *lcr1* and *lcr1-C* cells were acclimated for 16h in LL (15 µmol photons m^-2^ s^-1^). After sampling for the LL conditions, light intensity was increased to 300 µmol photons m^-2^ s^-1^ (HL); samples were taken 1 h (RNA) or 4 h (protein and photosynthetic measurements) after exposure to HL. Shown are relative expression levels of qE-related genes at the indicated conditions normalized to WT LL (*n* = 3 biological samples, mean ± sd). **C**. Immunoblot analyses of LHCSR1, LHCSR3, PSBS and ATPB (loading control) of one of the three biological replicates, under the indicated conditions. **D.** Quantification of immunoblot data of all replicates in panel **c** after normalization to ATPB. Shown are the HL treated samples; WT protein levels were set as 1. **E.** NPQ and calculated qE (as an inset) 4h after exposure to HL (*n* = 3 biological samples, mean ± s.d). The p-values for the comparisons are based on ANOVA Dunnett’s multiple comparisons test and as indicated in the graphs (*, P < 0.005, **, P < 0.01, ***, P < 0.001, ****, P < 0.0001). Statistical analyses for panels **b** and **d** were applied on log10- transformed values.

Furthermore, we revisited the role of LCR1 in regulating CCM genes^17^ by analyzing expression of selected CCM genes in WT, *lcr1* and *lcr1-C* cells shifted from LL or darkness to HL, conditions favoring CCM gene expression^7^. We first confirmed that under our experimental conditions *lcr1* could not fully induce *LCI1* (**Supplementary Fig. 8a, b**) in accordance to the report of the discovery of LCR1^17^. Our analyses further showed a statistically significant impairment of *lcr1* in inducing genes encoding the Ci transporters LOW-CO2-INDUCIBLE PROTEIN A (LCIA), HIGH-LIGHT ACTIVATED 3 (HLA3) and BESTROPHINE-LIKE PROTEIN 1 (BST1) as well as the carbonic anhydrase CAH4, when shifted from LL or dark to HL (**Supplementary Fig. 8a, b**), indicating that the role of LCR1 in low-CO2 gene expression extends beyond the regulation of gene expression of *CAH1*, *LCI1* and *LCI6*^17^.

### PHOT-specific GRN reveals a novel repressor of qE

The light-dependent induction of LHCSR3 is predominantly mediated by the blue light photoreceptor PHOT^18^. To analyze the PHOT-dependent transcriptional regulators, we first inferred a GRN based solely on the RNAseq from samples of *phot* and WT acclimated to LL and after 1h exposure to HL (data set PH, **Supplementary Table 1**) using only the GENIE3 approach, reported to show good performance^46, 52^ and which is among the best performing approaches in our consensus network. Since the PH experiment contains a low number of samples (12 samples, 4 conditions) the inferred GRN will inevitably suffer from the effects of low statistical power and high collinearity. To mitigate these effects, we decided to only include interactions that are also present in the benchmarked consensus network. By determining the intersection of the two networks, we obtained a GRN that resolves regulatory interactions underlying the transcriptomic changes observed in the *phot* mutant, while borrowing the statistical power of the whole RNAseq compendium. We refer to the resulting network as PHOT-specific GRN (**Methods**, **Supplementary Table 5**).

When investigating the top 10 regulators of qE genes in this PHOT-specific GRN the interactions where weighted based on the importance score from GENIE3 (**Supplementary Table 6, Fig. 3a**). Among the interactions in this list of PHOT-dependent regulators, we recovered two known regulators of qE, namely, ROC75^28^ and CrCO^26, 27^(**Fig. 3a**). These observations are in line with an existing hypothesis^24^ suggesting that a CUL4-dependent E3-ligase targeting CrCO^,26^ acts downstream of PHOT. ROC75 has been previously reported to act independently of the PHOT signal based on qPCR studies of the mutant grown synchronously under different light spectra^28^. In our RNAseq data, gathered under continuous white light, we observed a significant difference in expression levels of ROC75 between WT and *phot* (log2 fold-change = 1.03, adj. p-value = 1.80*10^-7^).

**Fig. 3:**
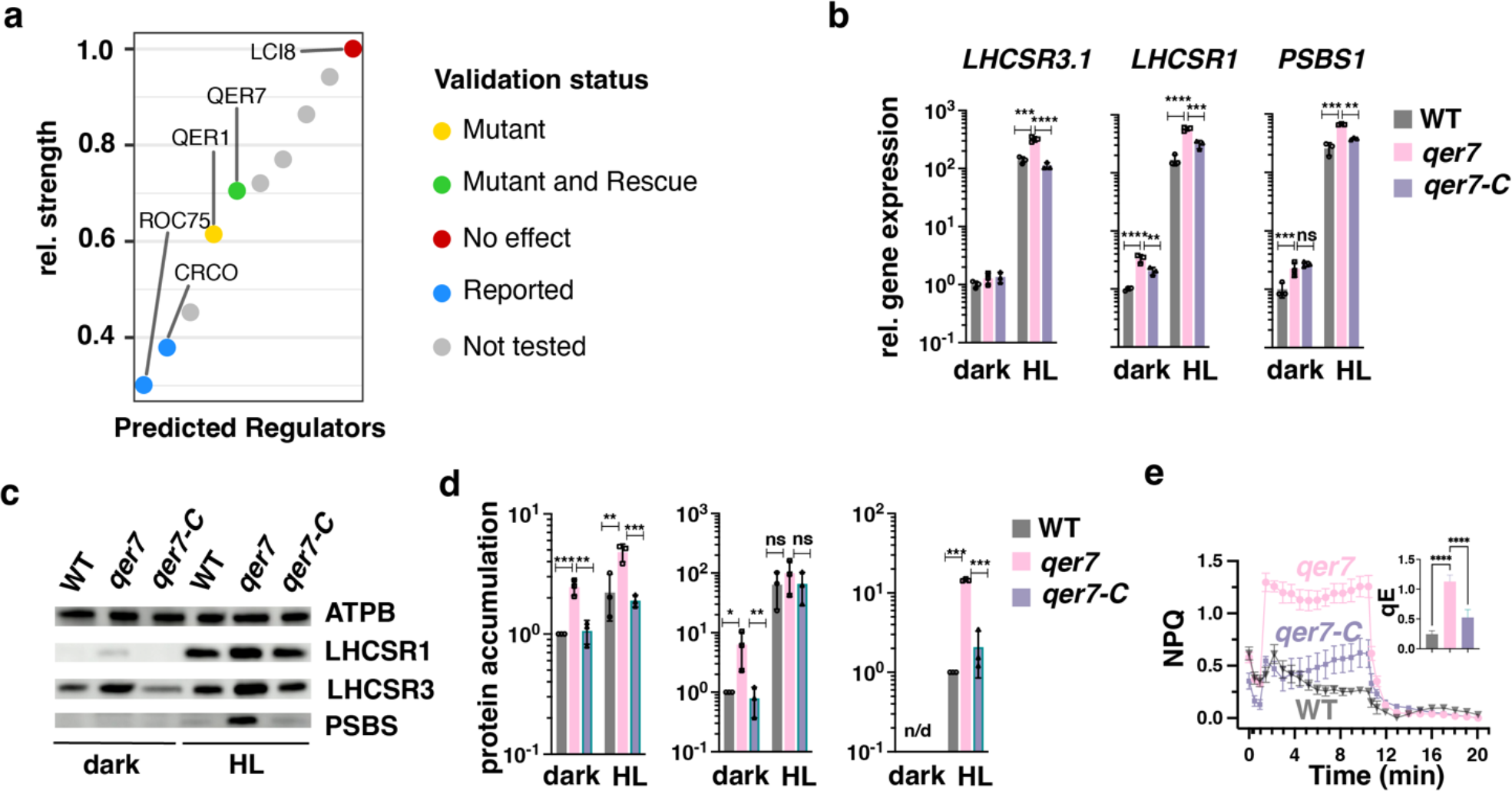
A PHOT-specific GRN pinpoints QER7 as a suppressor of the expression of qE-related genes. **A.** Dot plot of the relative regulatory strength of the top 10 regulators of qE-related genes in the PHOT-specific GRN (see **Methods**). TFs are marked in green if qE transcript levels were affected in the respective knock-out strain and this effect was reversed by complementation with the knocked-out gene, in yellow, if the effect was not reversed by complementation, and in red, if no mutant effect was observed in the mutant. TFs for which no mutant lines were available are plotted in grey. **B**. WT, *qer7* and *qer7-C* cells were synchronized under 12h light (15 µmol m^-2^ s^-1^)/12h dark cycles. After sampling for the dark conditions (end of the dark phase), cells were exposed to 300 µmol photons m^-2^ s^-1^ (HL); samples were taken 1 h (RNA) or 4 h (protein and photosynthetic measurements) after exposure to HL. Shown are relative expression levels of qE-related genes at the indicated conditions normalized to WT LL (*n* = 3 biological samples, mean ± sd). **C**. Immunoblot analyses of LHCSR1, LHCSR3 and ATPB (loading control) of one of the three biological replicate set of samples, under the indicated conditions. **D.** Quantification of immunoblot data of all replicates in panel **c** after normalization to ATPB. WT protein levels at HL were set as 1. **E.** NPQ and qE, measured 4h after exposure to HL (*n* = 3 biological samples, mean ± sd). The p-values for the comparisons are based on ANOVA Dunnett’s multiple comparisons test and are indicated in the graphs (*, P < 0.005, **, P < 0.01, ***, P < 0.001, ****, P < 0.0001). Statistical analyses for panel **b** and **d** were applied on log10- transformed values.

The fact that several regulators showed larger regulatory strength than CrCO in the PHOT-specific GRN indicates the existence of yet unreported regulators of qE effector genes in the PHOT signaling pathway. This is in line with existing results^26^, showing that the knock-out of CrCO is insufficient to fully abolish light-dependent activation of LHCSR3. Following this reasoning we obtained *qer1* and *qer7*, the available regulator candidates mutants, from the CLiP library^49^ (for genotyping see **Supplementary Fig. 5**). Our results show higher mRNA levels of LHCSR1 in the *qer1* mutant (**Supplementary Fig. 9**); however, this could not be rescued by ectopic expression of the *QER1* gene in the *qer1* mutant background (**Supplementary Fig. 9**). We found significant upregulation of *LHCSR3.1* gene expression in the *qer7* mutant (1.7 times, **Supplementary Fig. 10a**) also reflected in higher NPQ (**Supplementary Fig. 10b)** and qE levels (**Supplementary Fig. 10c**) which we followed up in more detail. To this end, we ectopically expressed the WT *QER7* gene in the *qer7* mutant and generated the complemented strain *qer7-C* that expressed *QER7* to levels similar to those WT (**Supplementary Fig. 5c**). As a result, the *qer7-C* strain showed reduced *LHCSR3* gene expression, NPQ and qE levels as compared with the *qer7* mutant (**Supplementary Fig. 10a-c**). *LHCSR1* and *PSBS* seemed to be unaffected in the qer7 in these LL to HL transition experiments (**Supplementary Fig. 10a**). As with LCR1, we also performed dark to HL experiments to further characterize the photoprotective responses of *qer7*; under these conditions, *qer7* accumulated significantly more *LHCSR1* (1.7 times) and *PSBS1* (2.2 times) while *LHCSR3* remained unaffected (**Supplementary Fig. 10d**). As in the LL to HL experiments (**Supplementary Fig. 10b, c**) *qer7* showed more NPQ and qE (**Supplementary Fig. 10e, f**). Complementation of *qer7* with the missing *QER7* gene restored all phenotypes (*LHCSR1*, *PSBS*, NPQ, qE; **Supplementary Fig. 10d-f**). These data validated the prediction of QER7 as regulator of qE gene expression (**Fig. 3a**) and indicated that QER7 regulates different subsets of qE genes depending on the pre-acclimation conditions; *LHCSR3* when preacclimated under LL, *LHCSR1* and *PSBS* when pre-acclimation occurs in darkness.

Motivated by these findings and given the fact that most of the Chlamydomonas transcriptome undergoes diurnal changes^39^ we decided to address the role of QER7 in regulating qE genes under light/dark cycles. We synchronized WT, *qer7* and *qer7-C* cells in 12h L/12h D cycle and exposed them to HL right after the end of the dark phase. Our results revealed that under these conditions QER7 functions as a repressor of all qE-related genes; the *qer7* mutant expresses significantly higher LHCSR3, LHCSR1 and PSBS not only at the gene (**Fig. 3b**) but also at the protein (**Fig. 3c, d**) level, and exhibits higher NPQ and qE (**Fig. 3e**), with all phenotypes rescued in the *qer7-C* complemented line. Previous protein homology studies identified QER7 as Squamosa Binding Protein^53^ or bZIP TF^54^, and here, we provide the first functional annotation of QER7 as a novel qE regulator.

### QER7 co-regulates qE-related and CCM genes

Our findings that the regulatory role of CIA5^7^ and LCR1 (**Fig. 2 and Supplementary Fig. 7**) extends beyond CCM to also control qE-related gene expression, prompted us to also inspect the expression levels of CCM genes in synchronized *qer7* cells (**Fig. 4a**). Indeed, for five of these transcripts (*HLA3*, *CAH4*, *BST1*, *LCI1*, *LCIA*) we observed a significant upregulation in *qer7* after HL exposure that was reversed by complementation with the *QER7* gene, indicating that QER7 suppresses expression of CCM genes; the suppression role of QER7 on CCM genes was only observable under HL, conditions that favor CCM gene expression^7^ and not in the dark (**Fig. 4a**). We subsequently checked if this regulation is also captured in the GRNs. To this end, we used the CCM genes included in **Fig. 4a** as target genes and predicted the top 10 regulators of these genes using the same method as for the qE regulators (**Methods**). Interestingly, we found LCR1 (**Supplementary Fig. 11a, c**) amongst the top regulators of both, CCM and qE genes, in the consensus network and QER7 in the PHOT- specific network (**Supplementary Fig. 11b, d)**.

**Fig. 4:**
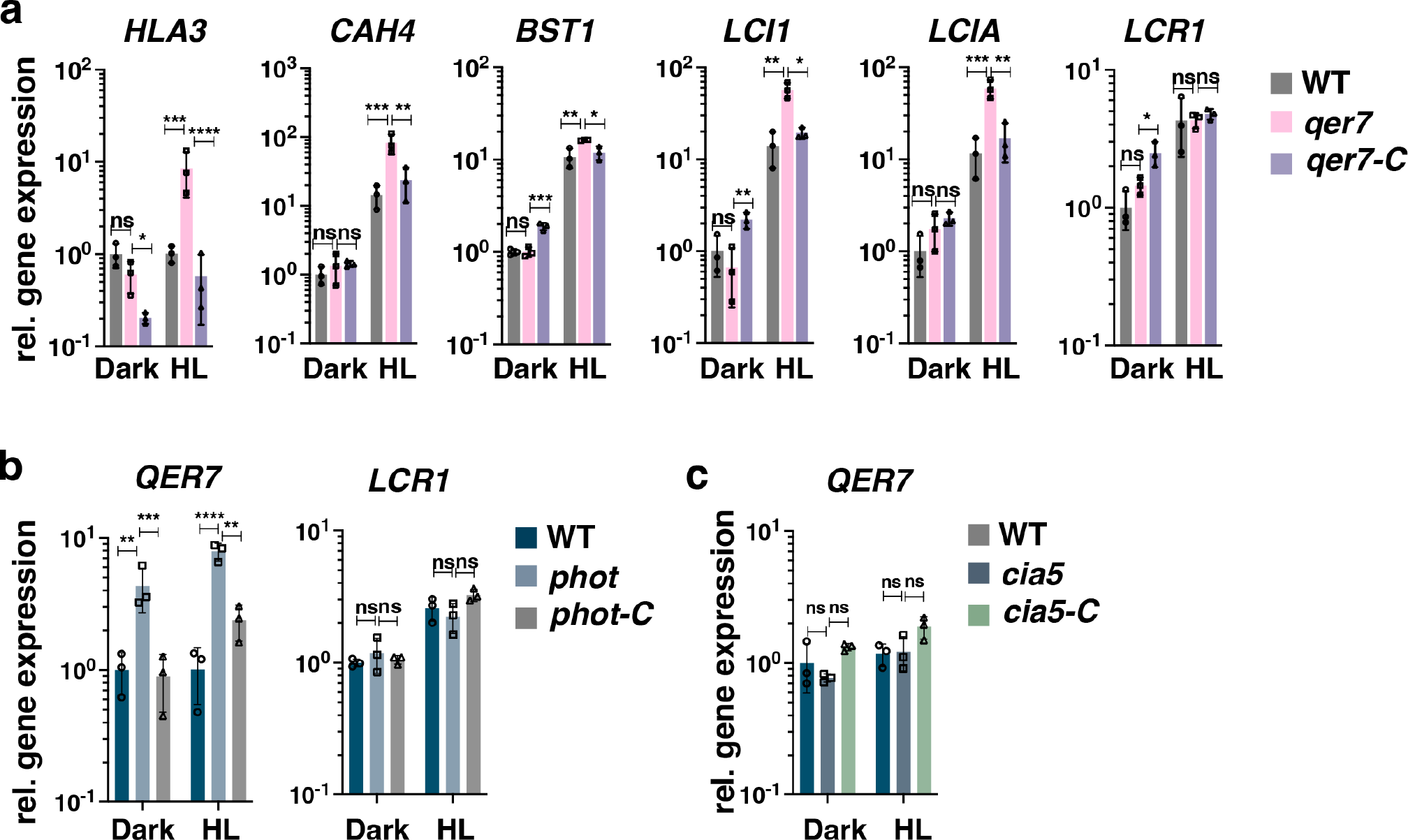
QER7 suppresses transcription of CCM genes and depends on PHOT. *Qer7*, *phot* and *cia5* cells alongside with their complemented lines (*qer7-C*, *phot-C* and *cia5-C* respectively*) and* WT backgrounds, were synchronized under 12h light (15 µmol m^-2^ s^-1^)/12h dark cycles, in photoautotrophic conditions (Sueoka’s high salt medium; HSM), at 23 °C in Erlenmeyer flasks shaken at 125 rpm. After sampling at the end of the dark phase, cells were exposed to 300 µmol photons m^-2^ s^-1^ (HL) and samples were taken 1 h after HL exposure. **A.** Relative expression levels of CCM genes at the indicated conditions. **B** Relative expression levels of *QER7* and *LCR1* in WT, *phot* and *phot-C*. **c** Relative expression levels of *QER7* in WT, *cia5* and *cia5-C.* In all cases (**a**, **b** and **c**) expression levels were normalized to WT LL (*n* = 3 biological samples, mean ± sd). The p-values for the comparisons are based on ANOVA employing Dunnett’s multiple comparisons test on log10-transformed values and are indicated in the graphs (*, P < 0.005, **, P < 0.01, ***, P < 0.001, ****, P < 0.0001).

Led by this observation, we investigated the signaling pathway upstream of QER7. To this end, we quantified *QER7* gene expression in synchronized *phot* cultures and observed that *QER7* is overexpressed in the *phot* mutant, suggesting that PHOT suppresses *QER7* expression (**Fig. 4b**). In contrast to *QER7*, *phot* expressed *LCR1* to levels similar to those in WT (**Fig. 4b**), and the same was true for the *qer7* mutant that also expressed *LCR1* to WT levels (**Fig. 4a**). Further validation that LCR1 and QER7 act on different pathways comes from the fact that although LCR1 is controlled by CIA5^55^, QER7 is not (**Fig. 4c)**. Thus, while sharing part of their target genes, the two TFs, LCR1 and QER7, mediate different signals. We captured this distinction in our PHOT-specific network, further underlining the power of the inferred networks.

Our findings that QER7 represses CCM gene expression (**Fig. 4a**) naturally raised the question whether the *qer7* mutant has altered CCM function, i.e. altered capacity of the cells to accumulate inorganic carbon (Ci). To address this question, we compared WT and *qer7* cells for their affinity for Ci, under conditions where CCM is not fully induced (LL) and after inducing CCM by acclimation to HL, which leads to mRNA accumulation of CCM-related genes (see for example **Fig. 4a** but also our previous study^7^). Under these conditions no difference could be observed between WT and *qer7* (**Supplementary Fig. 12, Supplementary Table 7**). Our data suggest that PHOT, by repressing *QER7* (**Fig. 4b**), is also involved in the regulation of CCM related gene expression. We investigated this further by quantifying CCM gene expression in WT, *phot* mutant and the complemented line *phot-C*, in the samples collected from the experiment presented in Fig. 4b, i.e. synchronized photoautotrophic cultures shifted to HL right after the end of the dark phase. We first analyzed expression of qE genes that was found as expected^18, 24, 25^ to be under control of PHOT (**Supplementary Fig. 13a**). We then analyzed expression of the five CCM genes found to be repressed by QER7 (**Fig. 4a**); out of those, *CAH4* and *HLA3* were down-regulated in the *phot* mutant exposed to HL after the end of the dark phase in synchronized cultures (**Supplementary Fig. 13b**) and this phenotype was fully rescued in the *phot-C* complemented line, suggesting a potential involvement of PHOT in regulating expression of CCM-related genes. Nevertheless, the affinity of *phot* for Ci was not different than this of WT (**Supplementary Fig. 14, Supplementary Table 7**), in line with previous work reporting CCM to be induced to very similar extent under blue or red illumination^56^. Thus, under our experimental conditions, the role of PHOT and QER7 in regulating CCM is restricted to the transcriptional level. Since the PHOT-QER7 pathway acts independently of the LCR1 pathway, both regulating the same subset of CCM genes we tested, it may be not so surprising that neither PHOT nor QER7 impact the affinity for Ci; in both *qer7* and *phot* mutants LCR1 levels are unaffected and therefore the control of LCR1 on CCM may mask any potential effect that PHOT or QER7 might have. Since many known CCM regulatory mechanisms act pots-transcriptionally^57^, it is conceivable, that the transcriptional regulation of PHOT and QER7 on their own are not sufficient and rely on integration with other simultaneous signals to clear all roadblocks for full CCM induction under HL.

### Genome-scale GRN indicates that photoprotection and CCM are co-regulated

The two qE regulators that we validated in this study also regulate CCM genes. Therefore, we next investigated to what extent the observed coregulation pattern applies to the global, known transcriptional regulation of low CO_2_ and light stress responsive genes. To this end, we took advantage of the size of the presented genome-scale GRNs and compiled a list of genes putatively involved in photoprotection (**Supplementary Table 8**) or the CCM (**Supplementary Table 9**); we then extracted the 10 TFs exhibiting the strongest regulatory strength on the genes in the compiled lists. We found six (empirical p-value<0.001, **Methods**) and four (empirical p-value < 0.01) of the top 10 regulators to be shared between these two responses in the consensus **(Fig. 5a, b, Supplementary Table 10**) and the PHOT-specific GRN, respectively (**Figure 5c,d, Supplementary Table 11**). The significant, large number of shared regulators is a strong indication that co-regulation of photoprotective and carbon assimilatory processes is a principal feature of Chlamydomonas’ transcriptional regulatory program.

**Fig. 5:**
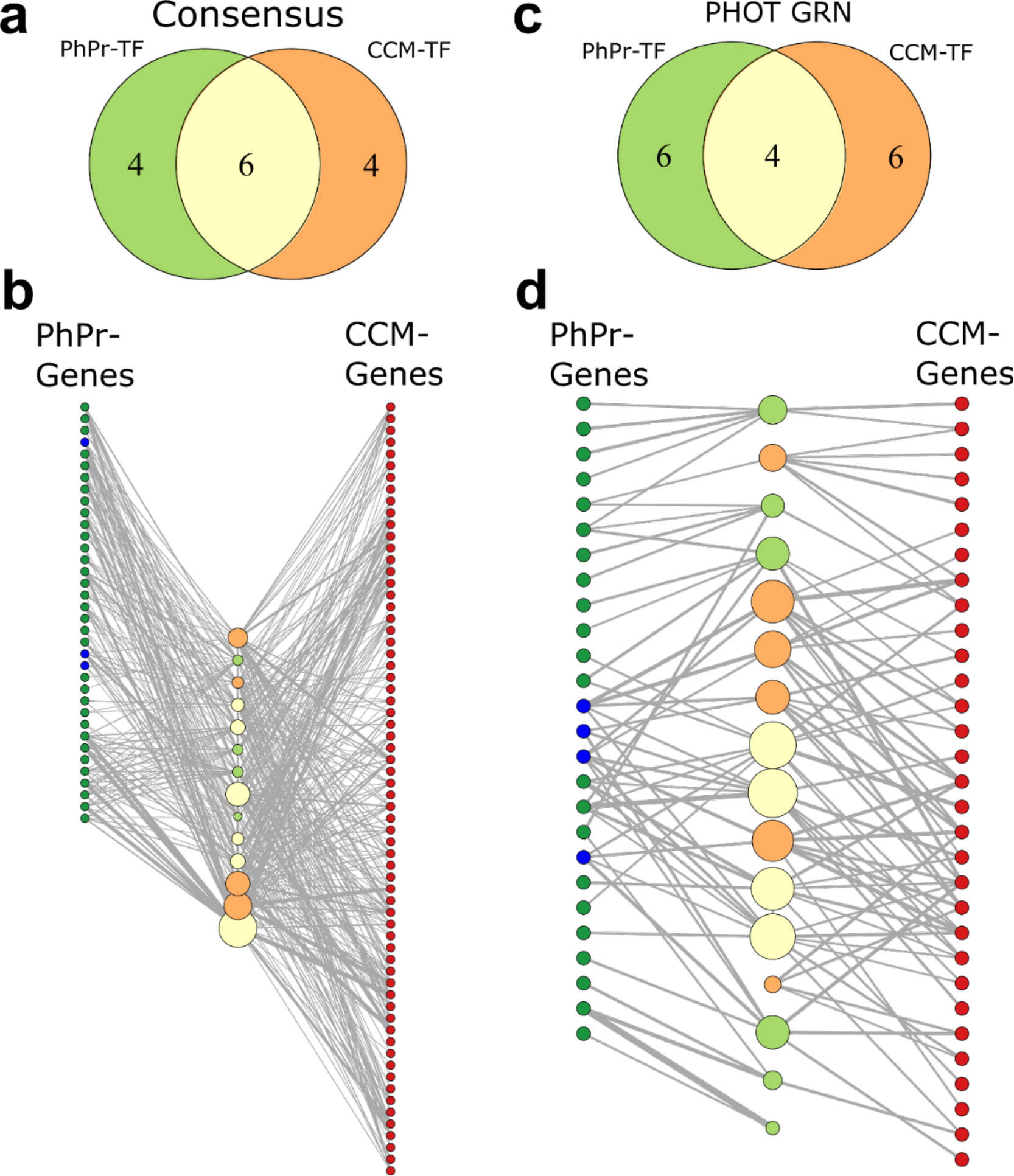
Consensus and PHOT-specific GRNs indicate extensive co-regulation of CCM and photoprotective genes. Top: Venn diagram depicting the overlap of the top 10 predicted TFs of the curated CCM and photoprotective (PhPr) genes based on **a.** the consensus or **c.** PHOT-specific GRN. Bottom: Network representation of the top ten TF sets of **b.** the consensus network or **d.** the PHOT-specific GRN (center nodes, same color code as in panel **a**) and the target genes used for prediction (left and right columns of nodes, photoprotection genes are shown in green, qE-related genes in blue, and CCM genes in red. In **b** the plotted regulatory strength corresponds to 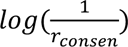, in **d**., it corresponds the GENIE3 edge weights denoting random forest importance measure. The edge width is proportional to the strength of the specific regulatory interaction. Size of the TF nodes corresponds to the sum of all plotted target gene edge weights.

## Discussion

The molecular actors and structure of the transcriptional regulatory mechanisms that shape *Chlamydomonas*’ response to differential light and carbon availability are largely unknown, although they are paramount to survival of *Chlamydomonas* and offer valuable targets for biological engineering. Here we set out to elucidate the GRN underlying the response to light and carbon availability by combining the results from five complementary inference approaches and data from 158 RNAseq samples of cultures responding to these cues. In the network inference process for this study, we carefully choose approaches and integration procedures developed to address the inherent difficulties of RNAseq data sets (e.g. collinearity and high number of variables compared to samples). Together with the many post-transcriptional layers of regulation in eukaryotic cells, the task of recovering the true GRN, nevertheless, remains a major challenge of the field of systems biology and this study is no exception.

We were able to experimentally validate two of the six novel qE regulators which the GRN predicted and for which KO mutants were available: QER7, suppressing LHCSR3, LHCSR1 and PSBS expression (**Fig. 3**, **Supplementary Fig. 10**), and LCR1, activating LHCSR3 and suppressing LHCSR1 and PSBS expression (**Fig. 2**, **Supplementary Fig. 7**). Both TFs also regulate expression of CCM genes, LCR1 as previously reported^17^, QER7 as demonstrated in this study (**Fig. 4**), suggesting that the processes of qE and CCM are co-regulated.

From the physiological point of view, the interconnection of qE and CCM comes as no surprise; exposure to HL boosts CO_2_ fixation rates and results in depletion of intracellular Ci, reflected, for instance, in the expression levels of the CO_2_-responsive marker RH1^7^; therefore, HL-exposed cells need to activate not only qE, to protect against photooxidative stress, but also CCM to sustain photosynthetic CO_2_ fixation. On the other hand, photooxidative damage is exacerbated by CO_2_-limitation; when CO_2_ fixation decreases, photosynthetically generated electrons accumulate in the electron chain potentially leading to reactive oxygen species generation^58^. Therefore, acclimation to low-CO_2_ availability needs to include not only activation of CCM to elevate CO_2_ levels at the site of fixation, but also of protection against photooxidative damage. It was recently shown that overexpression of the bZIP transcription factor BLZ8, resulting in enhanced CCM via overexpression of HLA3, CAH7 and CAH8, conferred enhanced oxidative tolerance triggered by alkaline stress ^59^. Although the qE capacity of the BZL8 overexpressing lines was not assessed, this work suggests that CCM and protection from oxidative stress are physiologically interconnected.

We complemented the findings of the involvement of LCR1 and QER7 in the regulation of qE and CCM related genes with an unbiased analysis of the genome-scale co-regulation of CCM and photoprotective genes. In this way we observed a significant number of regulators targeting both processes in the inferred consensus GRN as well as the PHOT-specific GRN (**Fig. 5**). This finding is in line with several experimental studies: *LHCSR3* mRNA has been reported to accumulate under low CO2^21, 22^ while exposure to HL has been reported to trigger CCM protein^60^ and mRNA^7^ accumulation. The Chloroplast Calcium Sensor protein CAS initially found to be a qE regulator^19^ turned out to also regulate CCM ^61^, and similarly, CIA5, the master regulator of CCM gene expression^14–16^ is also controlling qE gene and protein expression^7^. Our results indicate a set of TFs (**Fig 5, Supp. Table 10-11**) that likely act downstream of these different signals and integrate them into a common transcriptional response. Their further characterization is a promising avenue for future research.

We found QER7 in the list of PHOT-specific regulators and validated its role experimentally: Expression of *QER7* was repressed by PHOT (**Fig. 4b)** and was CIA5-independent (**Fig. 4c),** while expression of *LCR1* is regulated by CIA5^55^ and was found to be PHOT-independent (**Fig. 4b**). Thus, we showed that filtering the consensus network by differential expression data of mutants indeed successfully captures this context- specific regulatory interactions. As far as qE is concerned, qE genes are overexpressed in the *qer7* mutant (**Fig. 3b**), lacking the repressor QER7, and down-regulated in *phot* mutant (**Supplementary Fig. 13a**), overexpressing the QER7 repressor; as for CCM, QER7 represses expression of all five CCM genes we investigated (**Fig. 4a**), with two of them, *CAH4* and *HLA3*, being down-regulated in the *phot* mutant under HL (**Supplementary Fig. 13b**). Further work is needed to obtain a global understanding of the role of phototropin on the transcriptional regulation of Chlamydomonas CCM, including its observed role in controlling some of the CCM genes in the dark (**Supplementary Fig. 13b**); nevertheless, this suggested link forms an interesting parallel with the convergence of phototropin- and CO2-mediated signals recently shown to control stomata opening, responsible for CO2/O2 exchanges, in the model plant Arabidopsis ^62^.

In summary, we presented three valuable sets of PHOT-specific and general regulators: (i) a set of regulators of qE for which we validated available mutants (**Fig. 2a, Fig. 3a, Supp. Tables 4,6**), (ii) a set of regulators of the core CCM genes analysed via qPCR in **Fig. 4**, in which we recovered QER7 and LCR1 as coregulators of expression for qE and CCM genes (**Supplementary Fig. 11**), and (iii) a set of regulators **(Fig. 5, Suppl. Table 10-11**) of genes putatively involved in photoprotection or CCM (**Supp. Tables 8-9**), which depicted significant coregulation at a global scale.

Led by these predictions we experimentally showed that QER7 acts as a repressor of qE and CCM gene expression, LCR1 is a regulator of qE with a more expanded role on regulating CCM as previously thought and finally introduced a photoreceptor-mediated layer of regulation of CCM gene expression. These results clearly demonstrated that the generated GRN represents a powerful resource for future dissection of the transcriptional regulation of responses of *Chlamydomonas* to light and carbon availability. To allow easy access to this resource, we published an R-shiny webtool (https://github.com/arendma/GRN_web) to query the networks for arbitrary regulators and target genes. We expect that the webtool will prompt more concerted, community-wide efforts in resolving the interactions between other pathways that integrate different environmental cues in *Chlamydomonas*.

### Methods Transcriptome analysis

We assembled a compendium of RNAseq data (**Supplementary Table 1**) that capture regulation of light- dependent processes by combining in-house produced RNAseq measurements with publicly available data from two studies of densely sampled diurnal cultures of Chlamydomonas^38, 39^. For the samples in the acetate time-resolved experiment, adapter sequences were specifically trimmed from raw reads using BBduk^63^ (ktrim=r k=30 mink=12 minlen=50). Raw reads of the diurnal transcriptome study from Strenkert et al.^39^ were obtained from NCBI GEO database (GSE112394). Reads were aligned to the Chlamydomonas reference transcriptome^64^ available from JGI Phytozome (Assembly version 5) using RNA STAR aligner. The BAM files obtained from these measurements were analyzed using HTSeq-count^65^ (stranded=reverse) to create raw read count files. The raw read counts from Zones et al.^38^ were obtained as .tsv from NCBI GEO (GSE71469). The final data set consists of 158 samples from 62 experimental conditions or time points (**Supplementary Table 1**). Genes with less than 1 count per million in at least 9 measurements where discarded and the remainder were voom^64^ transformed and normalized using library normalization factors based on the TMM^66^ approach as implemented in the R Bioconductor package edgeR^67^.

### Transcription factor set from comparative genomics

To reduce the set of parameters in our network model, we compiled transcription factor (TF) annotations for the Chlamydomonas genome based on proteome homology studies. We obtained the proteomes and protein IDs of predicted Chlamydomonas TFs from Pérez-Rodríguez et al.^41^ Since these predictions were built based on the older Chlamydomonas assembly, we first used the conversion table provided by Phytozome to convert JGI4 to Crev5.6 IDs. For the TFs that could not be recovered by this approach we used the Phytozome BLAST tool to align these sequences against the Crev5.6 proteome (BLASTP, E threshold: -1, comparison matrix: BLOSUM62, word length: 11, number of reported alignments: 5). The reported hits were filtered for sequence identity > 97% and gaps ≤ 1 . If sequences mapped multiple times to the same Crev5.6 gene ID, only the hit alignment closest to the N-terminus of the query sequence was kept. The hit was only accepted if the alignment started at least six residues from the N-terminus of the hit sequence. For Crev5.6 loci that had multiple JGI4 TF queries assigned to them the best hit was selected manually. This set was then extended by the TFs found in the study of Jin et al.^42^ and the regulators in the manually curated set of CCM and qE regulatory interactions (**Supplementary Tables 8 and 9**). Using this procedure, we compiled a list of 407 Chlamydomonas TFs (**Supplementary Table 2**) to be considered as regulators in the inferred networks.

### Gene regulatory network inference

The CLR and ARACNE approach were based on all replicate measurements; for all other inference methods the median from each condition was used as input. All input matrices where standardized gene-wise. If not explicitly stated in the respective paragraph the implementations of all GRN inference approaches were applied with their default settings.

### Graphical Gaussian Models

The network inferred from a Graphical Gaussian model of gene regulation was obtained using the implementation of the partial correlation estimate from Schäfer et al.^34^ as implemented in the R GeneNet package. All interactions between TFs and another gene/TF with non-zero partial correlations were included as network edges.

### GENIE3

The random forest-based network from GENIE3 was generated using the R Bioconductor implementation provided by the authors^68^. We used only expression levels of TFs as predictors.

### Elastic net regression

A linear regression based network was obtained using the elastic net algorithm^35^. A model was fit for each gene using the expression levels of all TFs as predictors. The two hyperparameters λ2 (quadratic penalty) and s (fraction of L1 norm coefficients) were tuned for each gene model using 6-fold cross validation. The 2D parameter space scanned was λ2 = {0,0.001,0.01,0.05,0.1,0.5,1,1.5,2,10,100} and *s* = {0.1,0.2,0.3,0.4,0.5,0.6,0.7,0.8,0.9}. The R2 value for each model was calculated as

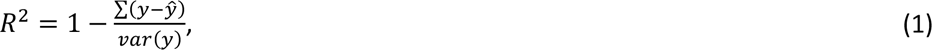

with *y* denoting the vector of observed expression values and *y*^ the model predictions. Models with a negative coefficient of determination (*R*2) value were discarded as regularization artifacts. The results of the remaining models were assembled into a network in which interactions were ranked by regression coefficients *β* normalized by the maximum absolute coefficient:

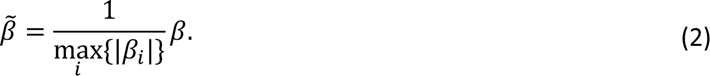

### CLR and ARACNE

The implementation of mutual information (MI)-based network inference approaches from the R package minet^69^ was used. Pairwise MI was estimated based on the Spearman correlation as proposed by Olsen et al.^70^. Two networks were constructed based on these MI estimates. Using the CLR approach^71^ non-significant interactions were removed based on the z-scores calculated from the marginal distributions of MI values for each gene pair. Alternatively, the ARACNE algorithm^44^ was used to prune the network based on the data processing inequality. For both networks only interactions originating from a TF were taken into consideration and edges were ranked according to the assigned MI value.

### Deconvolution and Silencing

For the two networks based on decomposition of the interaction matrix *G* the Pearson correlation matrix obtained from gene expression values was used as input.

The deconvolution approach introduced by Feizi and co-workers^72^ was implemented as previously described^46^.

The eigenvalue scaling factor *β* was initialized as *β*=0.9 and iteratively reduced in increments of 0.05 until the largest eigenvector of the direct interaction matrix generated by deconvolution was smaller than 1. Edges were ranked according to the deconvoluted interaction matrix.

The Silencing approach as described by Barzel et al.^45^ was implemented in R. The proposed approximation of the direct interaction matrix *S* in which spurious interactions are silenced relies on the invers of the observed correlation matrix *G*. In our implementation we used the Moore-Penrose pseudoinverse in case *G* was close to singular. In the resulting network edges were ranked according to the approximated silenced interaction matrix.

### Consensus network construction

To improve network quality^30^, we built a consensus network integrating the GRN models inferred by the different approaches introduced above. To this end, we used the Borda count election method^48^ whereby the rank *r* of an interaction *I* in the consensus network built on the predictions from *k* approaches is given by arithmetic mean of the ranks in the individual networks

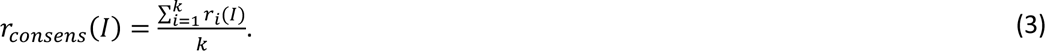

Following the reasoning of Feizi et al. ^72^ in this integration only the top 10% all possible edges in the GRN (625815) were considered from each individual ranking. For an edge that was not assigned a rank by some approaches, the missing ranks were set to 10% of all possible edges plus one.

Using this integration method, we assembled a consensus network based on all approaches to compare predictions from all GRN inference approaches (**Supplementary Fig. 4b**). Due to this comparison and their sensitivity of 0% (**Fig. 1b**) the rankings derived from ARACNE and Silencing were only considered in **Supplementary Fig. 4b** and excluded from the final consensus network used for all other analyses. As with the individual networks returned by the different approaches the consensus network **(Supplemental Table 3)** was trimmed to the top 10% of all possible edges according to the integrated ranks. For predictions of regulators the weight of edges in the final network was set as *r*^−^_*consens*_, denoting the reciprocal of the interaction rank.

### PHOT-specific network

To investigate the PHOT*-*specific regulatory interactions genes that are differentially expressed between phot mutant and wt under low and high light were inferred. To this end, transcript counts of genes with more than 1 count per million in at least four replicates from these conditions (**Supplementary Table 1**) were tested for differential expression using the R packages limma^73^, DeSeq2^74^, and edgeR^67^. Only genes deemed significant by all three tools after Benjamini-Hochberg correction for a false discovery rate of 0.05 were considered differentially expressed with respect to PHOT mutation In the next step, we focused on the normalized and scaled expression levels from these differentially expressed genes and the previously mentioned conditions, to infer a PHOT-specific GRN using GENIE3. To improve robustness of this network, which was obtained from a comparably sparse data set, we only considered the edges in the intersection with the final consensus network. Again, for both networks only the top 10% of possible edges were taken into account. Therefore, the obtained PHOT-network represents a subnetwork of the final consensus in which edges are weighted by the PHOT-specific GENIE3 importance measure **(Supplementary Table 5)**.

### Identification of major regulators

We compiled a manually curated list of possible target genes known to be involved in the processes of qE (*LHCSR1, LHCSR3.1/2, PSBS1/2*), photoprotection (**Supplementary Table 8**), and CCM (**Supplementary Table 9**). Based on the assumption that major regulators act on several genes important for a biological process, the regulatory strength of a candidate regulator (for the given process) was determined by the sum of edge weights *W*_*ij*_ between this regulator and the *k* genes in the respective target gene set

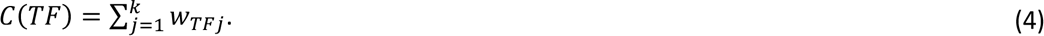

### Empirical p-value calculation using Monte-Carlo simulation

The one-sided p-value for the overlap between the regulators of CCM and photoprotective genes was approximated by sampling the overlaps of random gene sets. To this end, we compiled two gene sets with the same cardinality as the curated CCM and photoprotective genes. The genes in these sets where randomly sampled without replacement from all targets in the respective networks. The 10 strongest regulators of these two gene sets where then obtained as previously described and the overlap was calculated as our sample statistic. This process was repeated 10,000 times and an empirical p-value was calculated from the number of iterations, *r*, where the overlap was higher or equal to the observed value, and the total number of iterations, *n* ^75^:

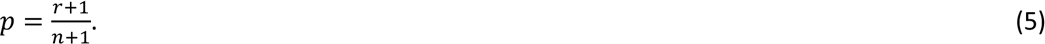

### Strains and conditions

*C. reinhardtii* strains were grown under 15 µmol photons m^-2^ s^-1^) in Tris-acetate-phosphate (TAP) media^76^ at 23 °C in Erlenmeyer flasks shaken at 125 rpm. For all experiments cells were transferred to Sueoka’s high salt medium (HSM)^77^ at 1 million cells mL^-1^ and exposed to light intensities as described in the text and figure legends. For the investigation of the impact of acetate on the genome-wide transcriptome, HSM was supplemented with 20 mM sodium acetate. *C. reinhardtii* strain CC-125 mt+ was used as WT. The *phot* mutant (depleted from *PHOT1*; gene ID: Cre03.g199000)*, was* previously generated^78^ and recently characterized together with its complemented line *phot-C*^25^. The *cia5* mutant (defective in *CIA5*, aka CCM1; geneID: Cre02.g096300; Chlamydomonas Resource Centre strain CC-2702), was previously generated^14^ and was used along with its complemented *cia5-C* (ref^7^). For synchronized cultures, the cells were grown in HSM for at least 5 days under a 12h light/12h dark cycle (light intensity was set at 15 μmol photons m^-2^ s^-1^; temperature was 18 ℃ in the dark and 23 ℃ in the light). All CLiP mutant strains used in this study and their parental strain (CC-4533) were obtained from the CLiP library (REF); *qer1* (LMJ.RY0402.072278), *qer4* (LMJ.RY0402.202963), *qer6* (LMJ.RY0402.162350), *qer7* (LMJ.RY0402.118995). The *lcr1* (strain C44)*, lcr1-C* (strain C44-B7) and its parental strain Q30P3 as described in^17^ were a kind gift from Hideya Fukuzawa. Before performing phenotyping experiments, we first confirmed that *lcr1* shows no expression of *LCR1* and that this is rescued in the *lcr1-C* strain (**Supplementary Fig. 15a**). The *lci8* overexpressing line was purchased from the Chlamydomonas Resource center; strain CSI_FC1G01, expressing pLM005-Cre02.g144800-Venus-3xFLAG in the CC-4533 background. Overexpression of LCI8-FLAG was verified by immunoblotting against FLAG (**Supplementary Fig. 15b**).

To complement *qer1*, a 1152 bp genomic DNA fragment from *Chlamydomonas* CC-4533 was amplified by PCR using KOD hot start DNA polymerase (Merck) and primers P11 and P12 (**Supplementary Table 12**). To complement *qer7*, a 5755 bp fragment DNA fragment from *Chlamydomonas* CC-4533 was amplified by PCR with Platinum superfii DNA Polymerase (Thermo Fisher Scientific) and primers P13 and P14 (**Supplementary Table 12**). The PCR products were gel purified and cloned into pRAM118^79^ by Gibson assembly^80^ for expression under control of the *PSAD* promoter. Junctions and insertion were sequenced and constructs were linearized by EcoRV before transformation. Eleven ng/kb of linearized plasmid^81^ mixed with 400 μL of 1.0 x10^7^ cells mL^-1^ were electroporated in a volume of 120 mL in a 2-mm-gap electro cuvette using a NEPA21 square-pulse electroporator (NEPAGENE, Japan). The electroporation parameters were set as follows: Poring Pulse (300V; 8 ms length; 50 ms interval; one pulse; 40% decay rate; + Polarity), Transfer Pluse (20V; 50 ms length; 50 ms interval; five pulses; 40% decay rate; +/- Polarity). Transformants were selected onto solid agar plates containing 20 μg/ml hygromycin and screened for fluorescence by using a Tecan fluorescence microplate reader (Tecan Group Ltd., Switzerland). Parameters used were as follows: YFP (excitation 515/12 nm and emission 550/12 nm) and chlorophyll (excitation 440/9 nm and 680/20 nm). Transformants showing high YFP/chlorophyll value were further analyzed by real time qPCR.

Unless otherwise stated, LL conditions corresponded to 15 µmol photons m^-2^ s^-1^ while HL conditions corresponded to 300 µmol photons m^-2^ s^-1^ of white light (Neptune L.E.D., France; see^7^ for light spectrum). All experiments were repeated at least three times to verify their reproducibility.

### DNA Isolation and genotyping of CLiP mutants

Total genomic DNA from CLiP mutants and corresponding wild-type strain CC-4533 was extracted according to the protocol suggested by CLiP website (https://www.chlamylibrary.org/). One μl of the extracted DNA was used as a template for the PCR assays, using Phire Plant Direct PCR polymerase (Thermo Fisher Scientific). To confirm the CIB1 insertion site in the CLiP mutants, gene-specific primers were used that anneal upstream and downstream of the predicted insertion site of the cassette (primer pairs P3-P4, P7-P8, P9-P10 and P5-P6 for *qer6*, *qer1*, *qer7* and *qer4* respectively; **Supplementary Table 12**). While all these primers worked in DNA extracted from WT, they did not work in the DNA extracted from the mutants, with the exception of *qer4* (**Supplementary Fig. 5**), therefore primers specific for the 5’ and 3’ ends of the CIB1 Cassette were additionally used. All the primers used for genotyping were shown in **Supplementary Table 12**. We further confirmed the disruption of the genes of interest by quantifying their mRNA accumulation (**Supplementary Fig. 5**).

### mRNA quantification

Total RNA was extracted using the RNeasy Mini Kit (Qiagen) and treated with the RNase-Free DNase Set (Qiagen). 1 μg total RNA was reverse transcribed with oligo dT using Sensifast cDNA Synthesis kit (Meridian Bioscience, USA). qPCR reactions were performed and quantified in a Bio-Rad CFX96 system using SsoAdvanced Universal SYBR Green Supermix (BioRad). The primers (0.3 μM) used for qPCR are listed in **Supplementary Table 13**. A gene encoding G protein subunit-like protein (GBLP)^82^ was used as the endogenous control, and relative expression values relative to *GBLP* were calculated from three biological replicates, each of which contained three technical replicates. All primers using for qPCR (**Supplementary Table 13**) were confirmed as having at least 90% amplification efficiency. In order to conform mRNA accumulation data to the distributional assumptions of ANOVA, i.e. the residuals should be normally distributed and variances should be equal among groups, Two-Way Analysis of Variance were computed with log-transformed data *Y* = log *X* where *X* is mRNA accumulation^83^.

### Immunoblotting

Protein samples of whole cell extracts (0.5 µg chlorophyll or 10 µg protein) were loaded on 4-20% SDS-PAGE gels (Mini-PROTEAN TGX Precast Protein Gels, Bio-Rad) and blotted onto nitrocellulose membranes. Antisera against LHCSR1 (AS14 2819), LHCSR3 (AS14 2766), ATPB (AS05 085), CAH4/5 (AS11 1737) were from Agrisera (Vännäs, Sweden); antiserum against PSBS was from ShineGene Molecular Biotech (Shanghai, China) targeting the peptides described in Ref.^9^. ATPB was used as a loading control. An anti-rabbit horseradish peroxidase–conjugated antiserum was used for detection. The blots were developed with ECL detection reagent, and images of the blots were obtained using a CCD imager (ChemiDoc MP System, Bio-Rad). For the densitometric quantification, data were normalized with ATPB.

### Fluorescence-based measurements

Fluorescence-based photosynthetic parameters were measured with a a pulse modulated amplitude fluorimeter (MAXI-IMAGING-PAM, HeinzWaltz GmbH, Germany). Prior to the onset of the measurements, cells were acclimated to darkness for 15 min. Chlorophyll fluorescence was recorded during 10 min under 570 µmol m^-2^ s^-1^ of actinic blue light followed by finishing with 10 min of measurements of fluorescence relaxation in the dark. A saturating pulse (200 msec) of blue light (6000 µmol photons m^-2^ sec^-1^) was applied for determination of Fm (the maximal fluorescence yield in dark-adapted state) or Fm’ (maximal fluorescence in any light-adapted state). NPQ was calculated as (*F*m – *F*mʹ)/*F*mʹ based on^84^; qE was estimated as the fraction of NPQ that is rapidly inducible in the light and reversible in the dark.

### CO2-dependent O2 evolution

The measurements were performed in accordance with ^85^ with minor modifications. Cells in photoautotrophic conditions (Sueoka’s high salt medium; HSM^77^), shaken in Erlenmeyer flasks at 125 rpm and 23 °C were shifted from LL (overnight at 15 µmol m^-2^ s^-1^) to HL (4h at 300 µmol m^-2^ s^-1^) in order to induce the CCM. Cells were suspended in 4ml 25 mM HEPES-KOH buffer (pH 7.3), at 25 µg chlorophyll per mL and were briefly sparged with nitrogen gas to remove the dissolved inorganic carbon (Ci). The cells were then transferred to an oxygen respiration vial (Pyro Science GmbH, Aachen, Germany) and were illuminated at 300 μmol photons m^-2^ s^-1^ for about 10-20 minutes, until no net oxygen evolution was seen, an indication that the internal Ci was depleted. Ci concentration in the cell suspension was then increased by stepwise injecting NaHCO3 with a microsyringe. O2 evolution was measured using a fiber optic probe (FireStingO2, Pyro Science GmbH, Aachen, Germany). Cumulative concentration of NaHCO3 after each addition were as follows: 25, 50, 100, 250, 500, 1000, 2000 μM. K1/2(Ci), the Ci concentration needed for half maximal rate of oxygen evolution, and Vmax were calculated by non-linear curve fitting to the Michaelis-Menten equation, using Prism Graph (GraphPad Software, LLC).

### Data and material availability

The consensus and PHOT-specific GRN are contained in edge list format as Supplemental Table 3 and Supplemental Table 5, respectively. To allow easy access to the information, we developed an R-shiny webtool that allows to query arbitrary TFs and target genes for regulatory interactions. The R-shiny webtool can be accessed at https://github.com/arendma/GRN_web. The code for GRN inference is available at https://github.com/arendma/GRN_code. The source data underlying Figures 2-4 and Supplementary Figures 2, 5c, 6, 7, 8, 9, 10, 12, 13, 14 and 15 are provided as a Source Data file. All biological material described in this study is available upon request.

## Acknowledgments

We are grateful to Prof. Hideya Fukuzawa for sending us *lcr1*, *lcr1-C* and their respective WT strains and to Prof. Peter Jahns for the antibody against PSBS.

## Supplementary Materials

### Supplementary Note 1

In the course of independent projects aiming to identify novel regulators of *LHCSR3*, we have investigated the role of four transcriptional factors, designated as TF1-4, in regulating expression of *LHCSR3.1* (*TF1*: MYB-like DNA-binding protein, Cre01.g034350; *TF2*: RWP8, RWP-RK transcription factor, Cre04.g218050; *TF3*: RWP5, RWP-RK transcription factor, Cre06.g285600; *TF4*: bHLH domain-containing protein, Cre07.g349152). Mutants bearing mutations in TF1-4 genes were ordered from the CliP library**^1^**. After confirming (Supplementary Fig. 2a) that expression of the genes of interest was abolished (*tf1-3* mutants) or was significantly higher than in WT (for the case of *tf4-oe*), we applied the following experimental setup to probe for *LHCSR3.1* expression: WT and mutant cells were acclimated for 16h in LL (15 µmol photons m^-2^ s^-1^). After sampling under the LL condition, light intensity was increased to 300 µmol photons m^-2^ s^-1^ (HL); samples for RNA extraction were taken after 1 h of exposure to HL. Our data showed that WT and mutants had similar expression levels of *LHCSR3.1* (**Supplementary Fig. 2**), excluding a role of these TFs in regulating transcription of *LHCSR3.1*.

### Supplementary Note 2

Simulation studies^2^ indicated that the top 10% of the edges in the consensus network are enriched in positive interactions if the underlying approaches perform better than guessing. Thus, a higher overlap with the consensus networks is expected to result in improved predictions. We found that the interactions from the network deconvolution approach exhibited the largest overlap with the consensus network, while GGM assigns the highest rank to consensus interactions (**Supplementary Fig. 4a**). Repeating this analysis by including the two worse performing approaches (i.e. ARACNE and Silencing) resulted in no qualitative changes in the overlap (**Supplementary Fig. 4b**) and only 7.47% of difference in the included interactions compared to the original consensus network, demonstrating the robustness of the inferred interactions. Considering the 10% network density threshold, we also inspected the proportion of TF-TF interactions in the different networks and their consensus; we found that it ranges from 2.55% in the GENIE3 network to 2.77% in the network deconvolution approach (**Supplementary Fig. 4c**). Since the fully connected network, containing all possible TF-TF and TF-target interactions, has a relative TF-TF interaction content of 2.52%, these findings suggest an enrichment of TF-TF interactions in GRNs inferred by all five approaches considered in the consensus.

---

**Supplementary Fig. 1:**
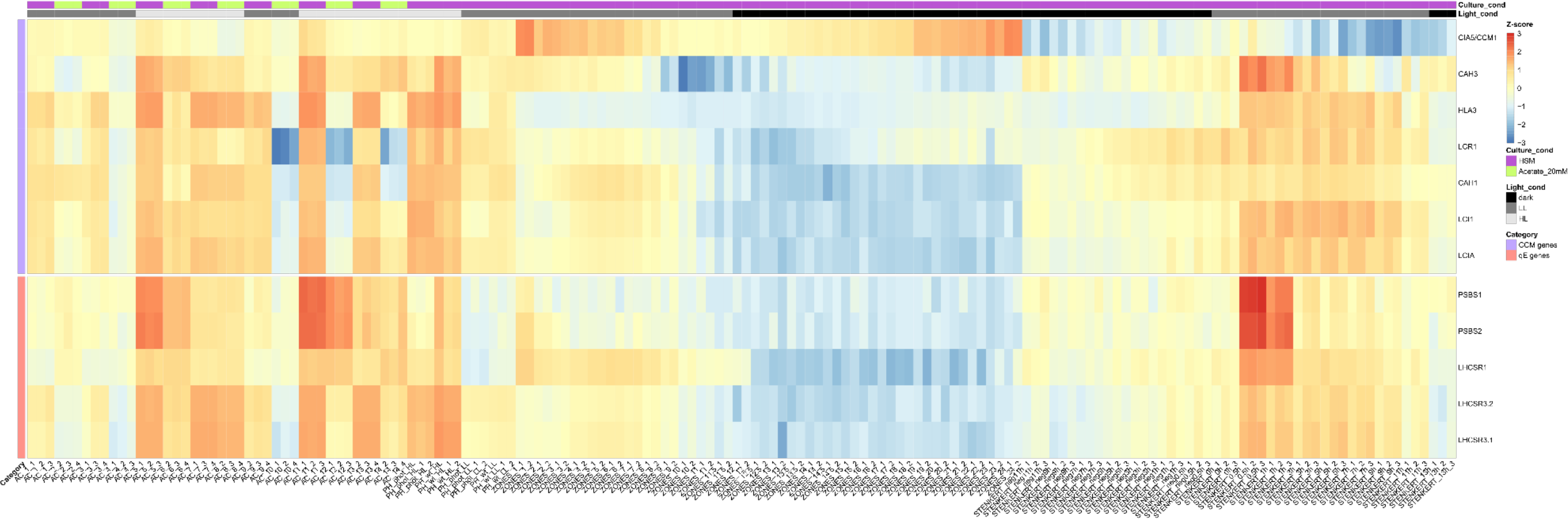
Z-score of log-transcript levels for representative CCM and qE genesin the used RNAseq data set. **Enlarged** representation of **Fig. 1a** with sample names included (see **Supplementary Table 1** for more details). The last digit of the column names indicates the replicate number. The column annotation provides information on the culture conditions. No clustering was applied.

**Supplementary Fig. 2:**
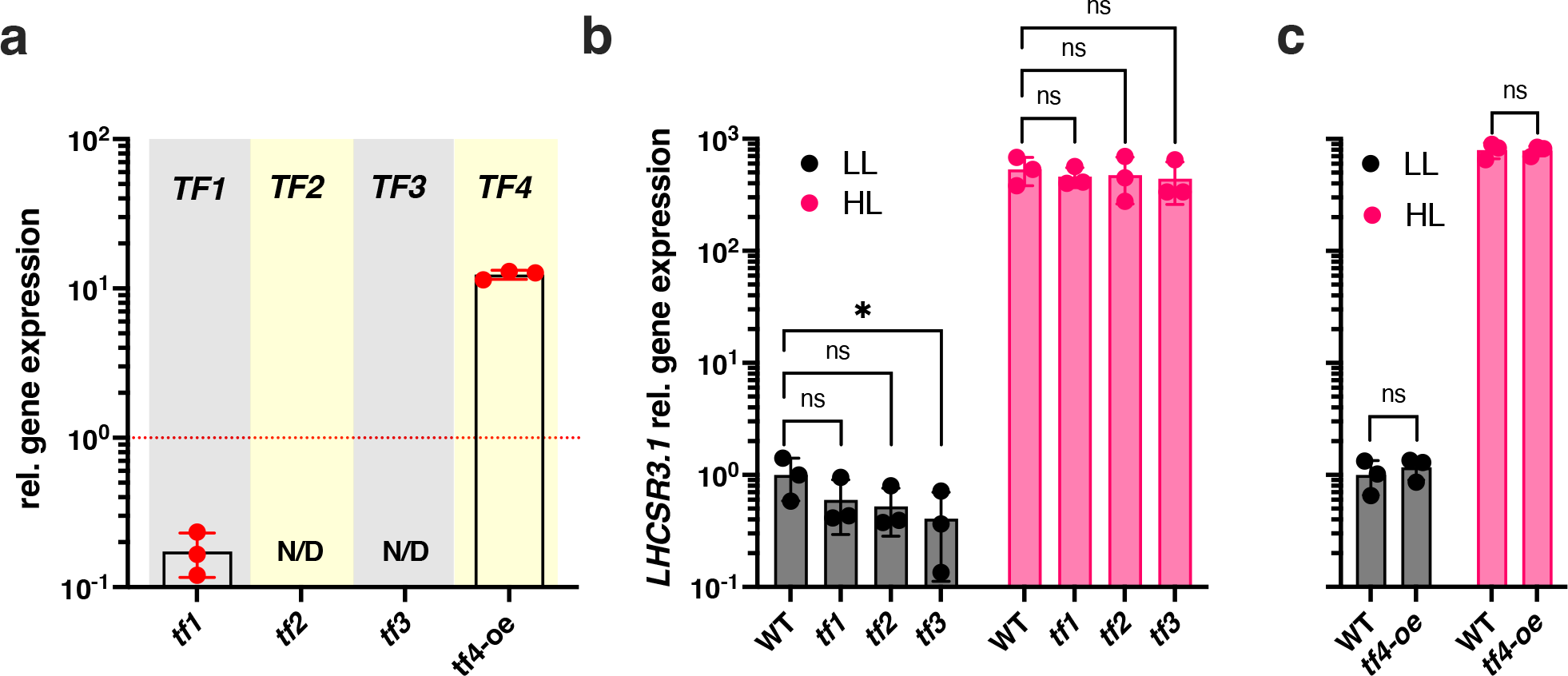
*LHCSR3.1* expression in mutants bearing mutations in transcription factors TF1-4. **a** Relative expression levels of *TF1-4 in tf1, tf2, tf3* and *tf4* mutant strains, grown under LL conditions in tris- acetate-phosphate (TAP) medium, normalized to WT (indicated as a red dashed line in the figure); N/D: non- detectable. **b** and **c** WT, *tf1, tf2, tf3 and tf4* cells were acclimated for 16h in LL (15 µmol photons m^-2^ s^-1^). After sampling under the LL conditions, light intensity was increased to 300 µmol photons m^-2^ s^-1^ (HL); samples were taken 1 h after exposure to HL. Shown are relative expression levels of *LHCSR3.1* at the indicated conditions normalized to WT LL (*n* = 3 biological samples, mean ± sd) for **b** WT (CC-4533) and tf1-3 and **c** for WT (CC-125) and tf4-oe, a strain overexpressing *tf4* in the CC-125 background. The p-values for the comparisons between the mutants and the WT are based on ANOVA Dunnett’s multiple comparisons test using log10- transformed values; the p-values are indicated in the graphs (*, P < 0.005; ns: non-significant). *TF1*: MYB-like DNA-binding protein, Cre01.g034350; *TF2*: RWP8, RWP-RK transcription factor, Cre04.g218050; *TF3*: RWP5, RWP-RK transcription factor, Cre06.g285600; *TF4*: bHLH domain-containing protein, Cre07.g349152).

**Supplementary Fig. 3:**
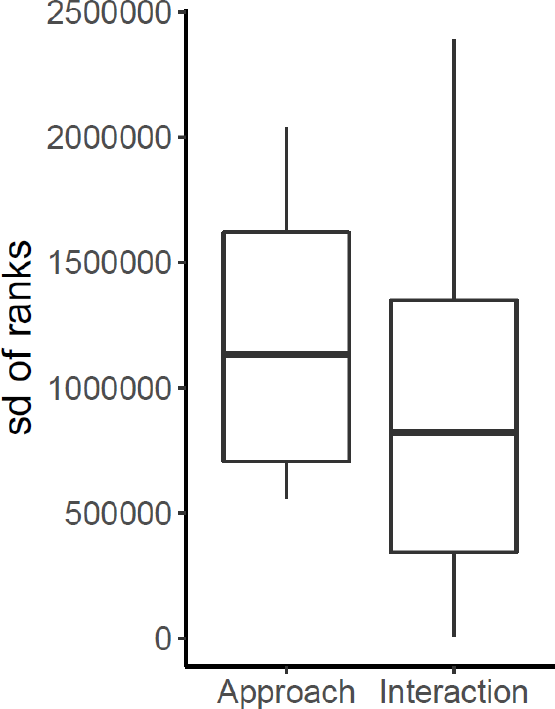
Variation in ranks of known interactions across approaches and interactions from different inference approaches. Boxplots of the standard deviation of ranks calculated for the curated interactions plotted in **Fig. 1b** over all ranks assigned by one approach or over all ranks assigned to one interaction.

**Supplementary Fig. 4:**
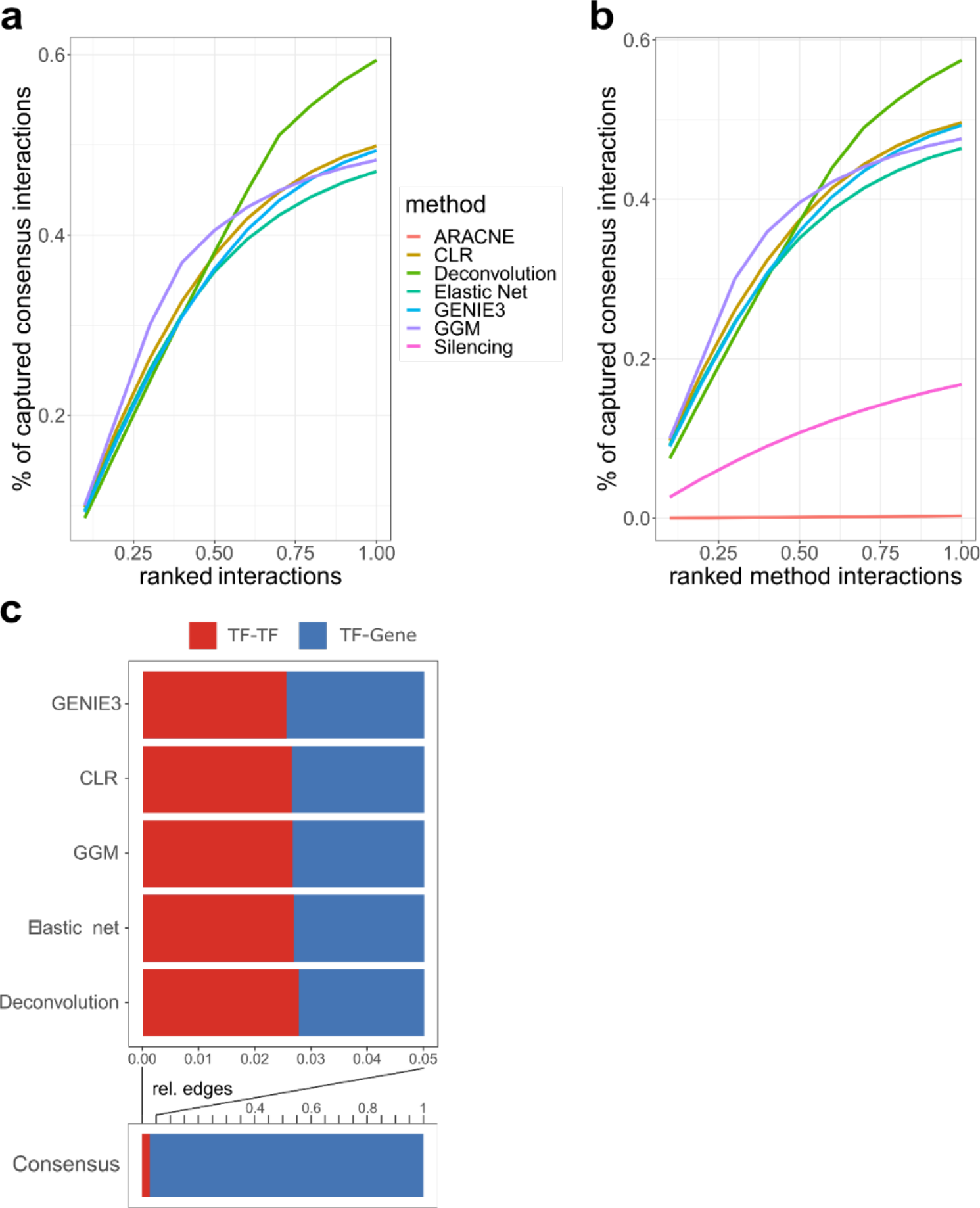
Overlap of consensus network with individual inference methods and proportion of TF-TF interactions in the obtained GRNs. **a.** The graph shows the overlap between the edges of the consensus network and the ranked interactions of the individual approaches normalized to the total number of edges in the consensus network used for regulator prediction. **b.** Same as in a but plotting the overlap with a consensus of all used approaches (including ARACNE and Silencing). **c**. Bar plots provide the proportion of TF-TF interactions in comparison to TF-target gene interactions contained in the GRNs inferred by the different approaches and the consensus GRN resulting from their integration. All depicted analyses considered only the interactions within in the 10% network density threshold (see **Methods**).

**Supplementary Fig. 5:**
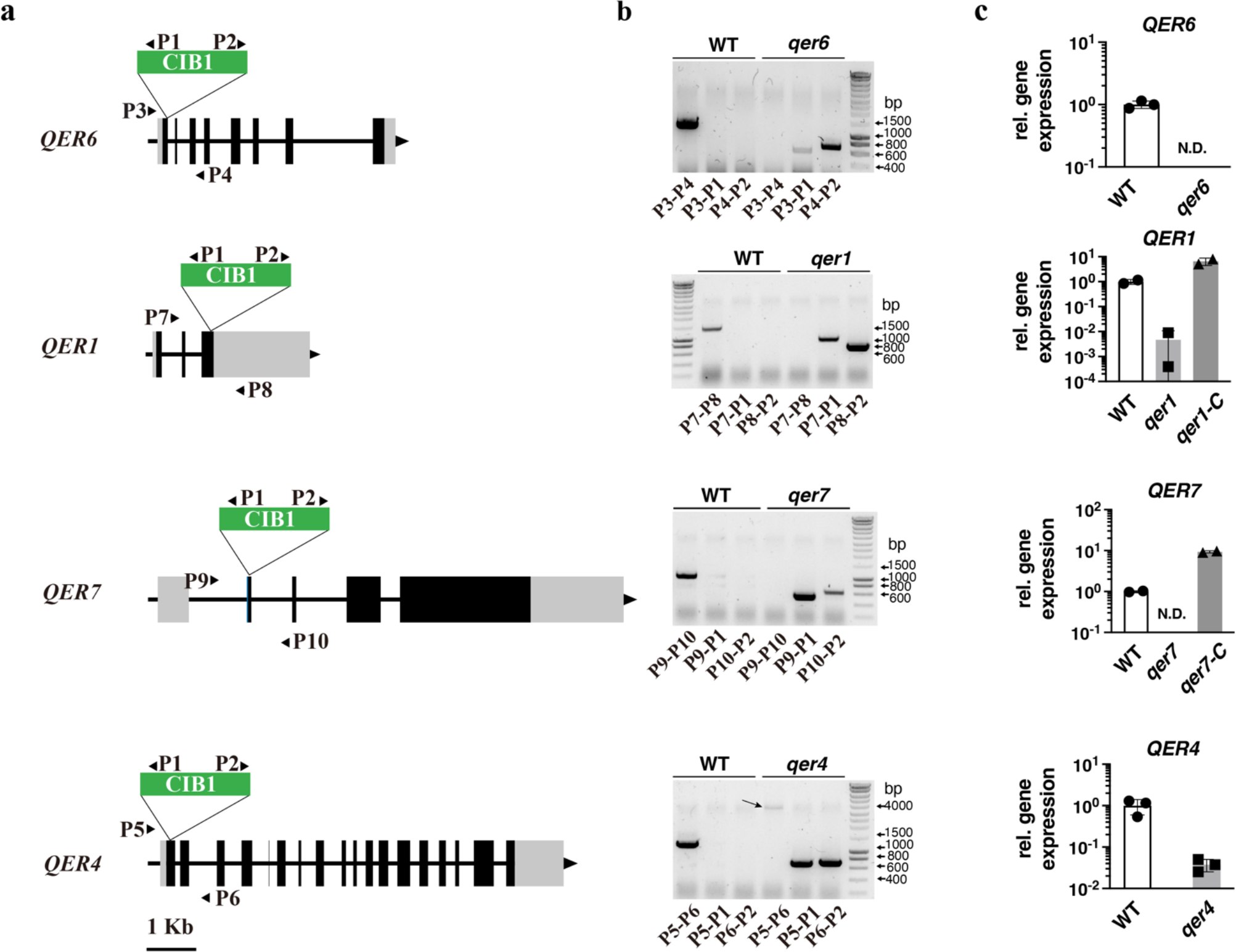
Genotyping of CLiP mutants affected in the predicted regulators of qE-related genes. **a.** Insertion map of the CIB1 cassette in the different genes. Exons are shown in black, introns as interconnecting lines, 5’UTR and 3’UTR in light gray, and primers in arrows. The insertion site of the CIB1 cassette is indicated by the triangle; **b**. PCR-validation of the insertion site in the different CLiP mutants using genomic DNA. To confirm the CIB1 insertion site, gene-specific primers were used that anneal upstream and downstream of the predicted insertion site of the cassette (primer pairs P3-P4, P7-P8, P9-P10 and P5-P6 for *qer6*, *qer1*, *qer7* and *qer4* respectively; **Supplementary Table 12**). Pairs of primers used are indicated at the bottom of the agarose gels used to separate the PCR products. Note that the PCR product of P5-P6 is indicated by an arrow. **c.** Relative expression levels of predicted qE regulator genes in the different CLiP mutants after exposure to HL (300 µmol photons m^-2^ s^-1^) for 1h.

**Supplementary Fig. 6:**
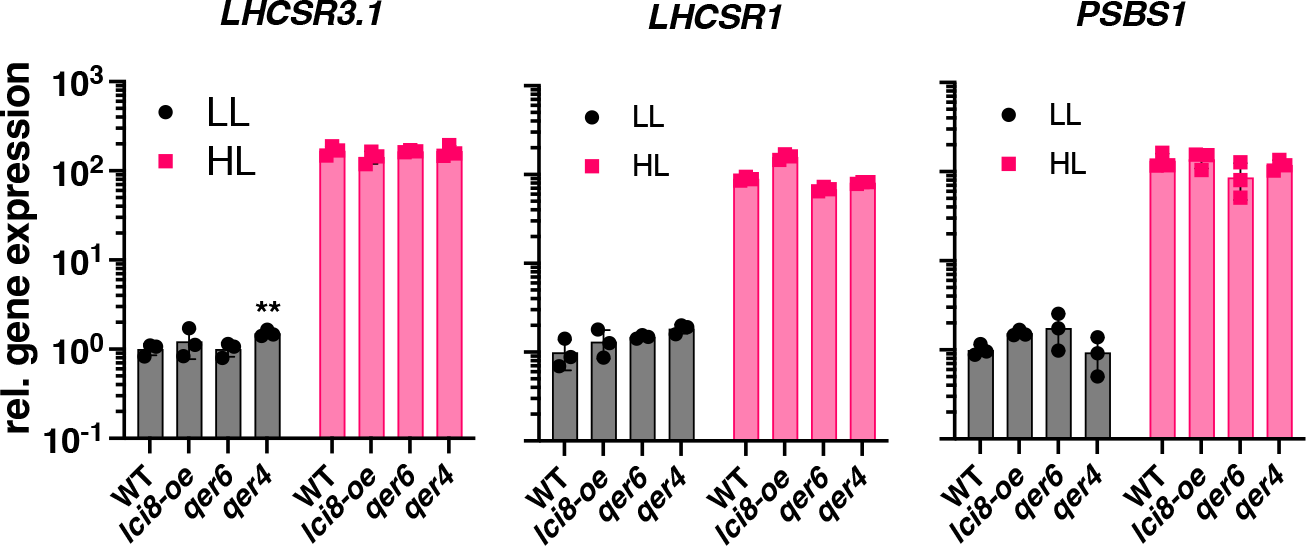
qE gene and protein expression in mutants bearing mutations in predicted qE regulator genes. WT, *lci8-oe, qer6* and *qer4* cells were acclimated for 16h in LL (15 µmol photons m^-2^ s^-1^). After sampling for the LL conditions, light intensity was increased to 300 µmol photons m^-2^ s^-1^ (HL); samples were taken 1 h after exposure to HL. Shown are relative expression levels of *LHCSR3.1*, *LHCSR1* and *PSBS1* at the indicated conditions normalized to WT LL (*n* = 3 biological samples, mean ± sd). The p-values for the comparisons between the mutants and the WT are based on ANOVA Dunnett’s multiple comparisons test on log10- transformed values and are indicated in the graphs (**, P < 0.01).

**Supplementary Fig. 7:**
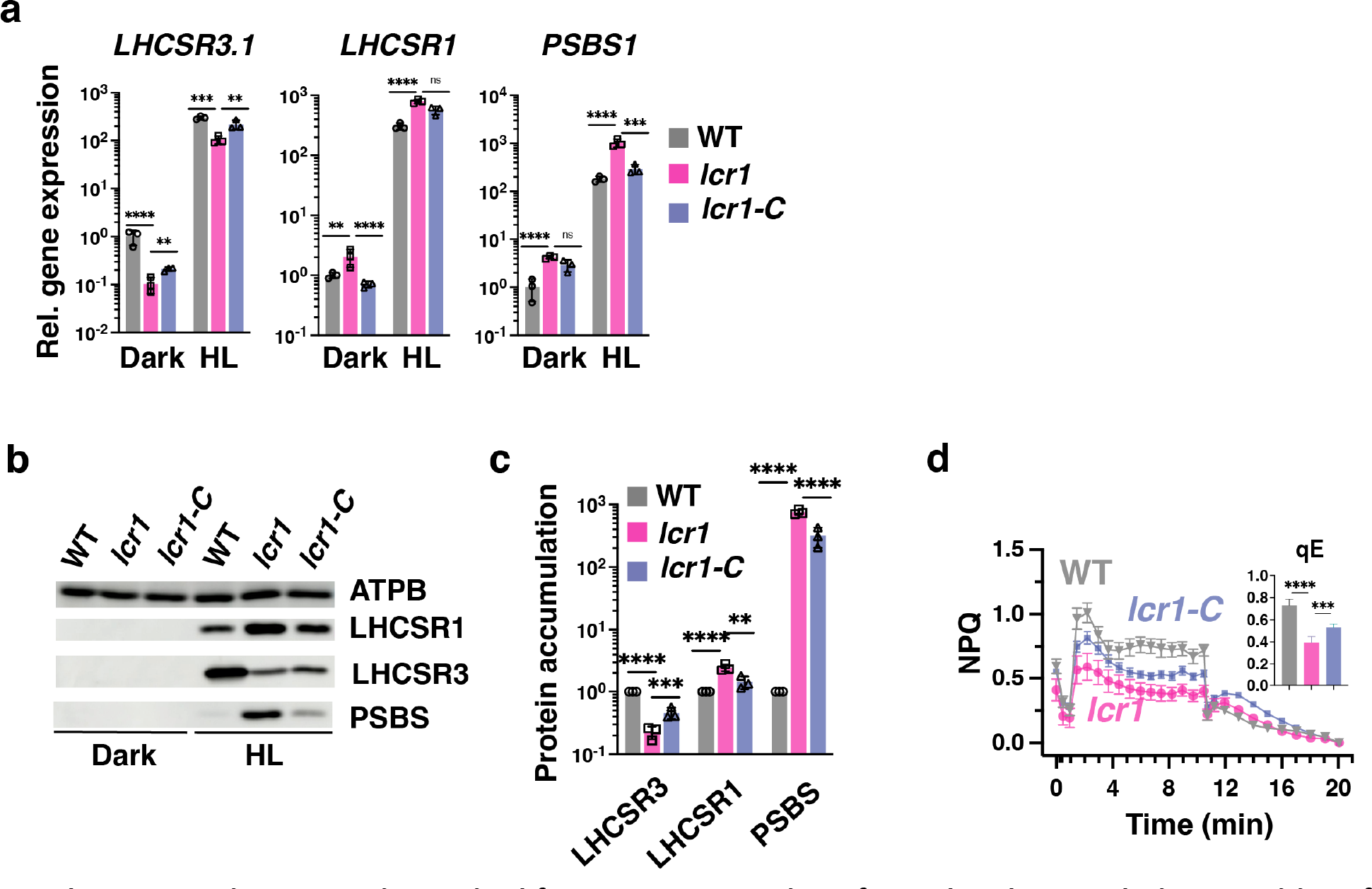
LCR1 is required for proper expression of qE-related genes during transitions from dark-to-light. WT, *lcr1* and *lcr1-C* cells were acclimated for 16h in darkness. After sampling for the dark conditions, light intensity was increased to 300 µmol photons m^-2^ s^-1^ (HL); samples were taken 1 h (RNA) or 4 h (protein and photosynthetic measurements) after exposure to HL. **a**. Relative expression levels of qE genes at the indicated conditions normalized to WT LL (*n* = 3 biological samples, mean ± sd). **b**. Immunoblot analyses of LHCSR1, LHCSR3, PSBS and ATPB (loading control) of one out of the three biological replicate samples, under the indicated conditions. **c.** Summary graph of immunoblots of all replicate samples of Supplementary Fig. 4b after normalization to ATPB. Shown are the HL treated samples; WT protein levels were set as 1. **d.** NPQ and calculated qE, 4h after exposure to HL (*n* = 3 biological samples, mean ± s.d). The p-values for the comparisons are based on ANOVA Dunnett’s multiple comparisons test and are indicated in the graphs (*, P < 0.005, **, P < 0.01, ***, P < 0.001, ****, P < 0.0001). Statistical analyses for panel **a** and **c** were applied on log10- transformed values.

**Supplementary Fig. 8:**
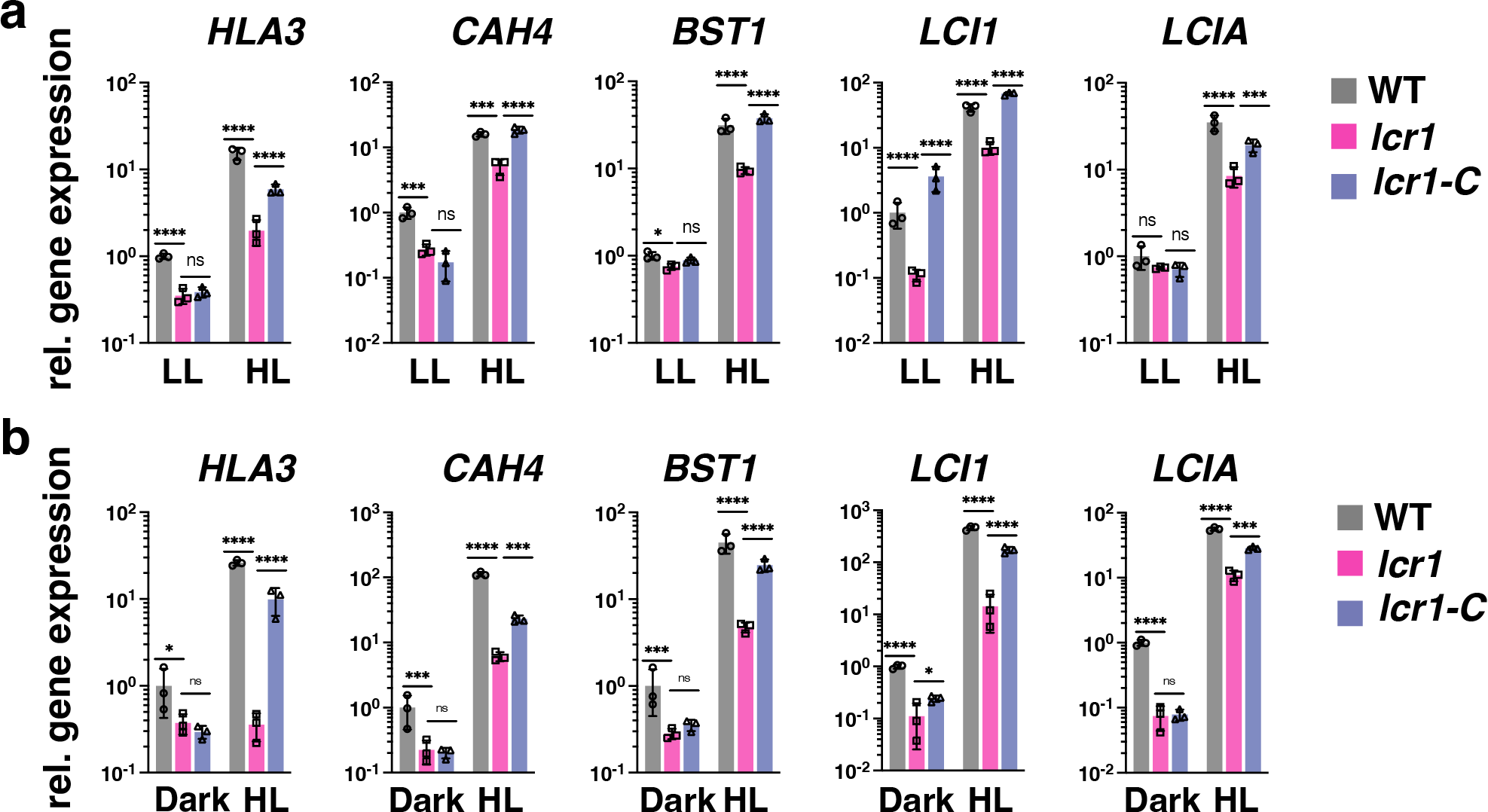
LCR1 activates the transcription of CCM genes in LL and dark-acclimated cells. **a.** WT, *lcr1* and *lcr1-C* cells were acclimated for 16h in LL (15 µmol photons m^-2^ s^-1^). After sampling under LL conditions, light intensity was increased to 300 µmol photons m^-2^ s^-1^ (HL) and samples were taken 1 h after exposure to HL. Relative expression of CCM genes at the indicated conditions normalized to WT under LL. **b**. Relative expression of CCM genes acclimated in 16h of darkness (*n* = 3 biological samples, mean ± sd). The p-values for the comparisons are based on ANOVA Dunnett’s multiple comparisons test on log10- transformed values and are indicated in the graphs (*, P < 0.005, **, P < 0.01, ***, P < 0.001, ****, P < 0.0001).

**Supplementary Fig. 9:**
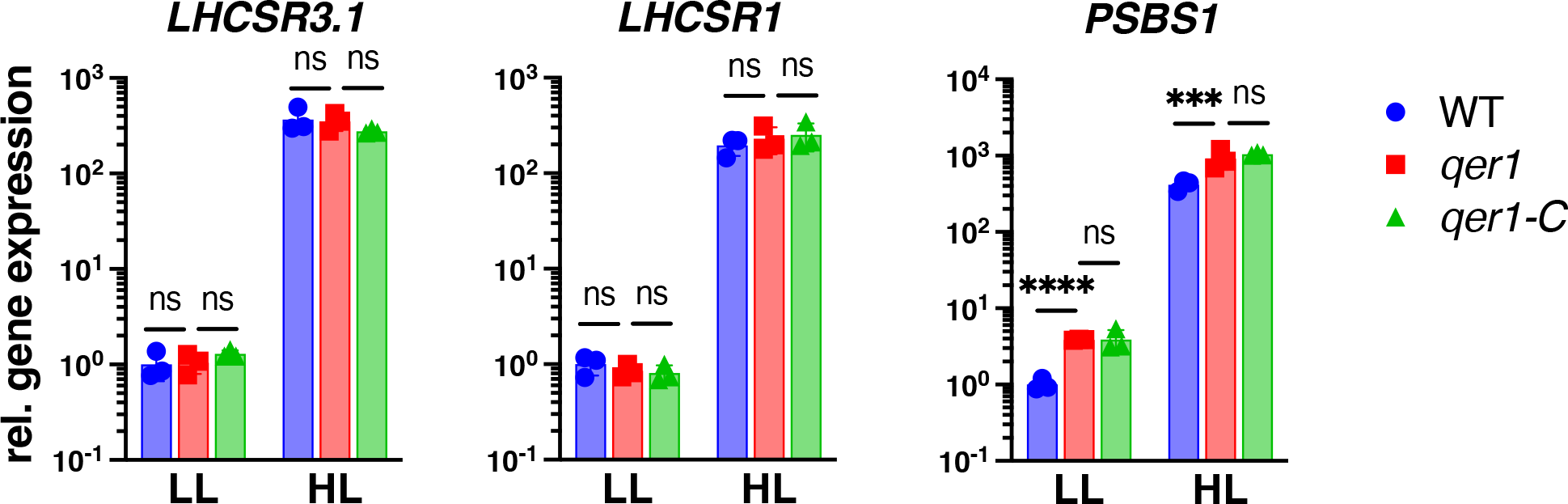
Complementation of *qer1* with *QER1* (*qer1-C* strain) fails to rescue the *LHCSR1* and *PSBS* phenotypes. WT, *qer1* and *qer1-C* cells were acclimated for 16h in LL (15 µmol photons m^-2^ s^-1^). After sampling for the dark conditions, light intensity was increased to 300 µmol photons m^-2^ s^-1^ (HL); RNA samples were taken 1 h after exposure to HL. Shown are relative expression levels of qE genes at the indicated conditions normalized to WT LL (*n* = 3 biological samples, mean ± sd). The p-values for the comparisons are based on ANOVA Dunnett’s multiple comparisons test on log10- transformed values and are indicated in the graphs (*, P < 0.005, **, P < 0.01, ***, P < 0.001, ****, P < 0.0001).

**Supplementary Fig. 10:**
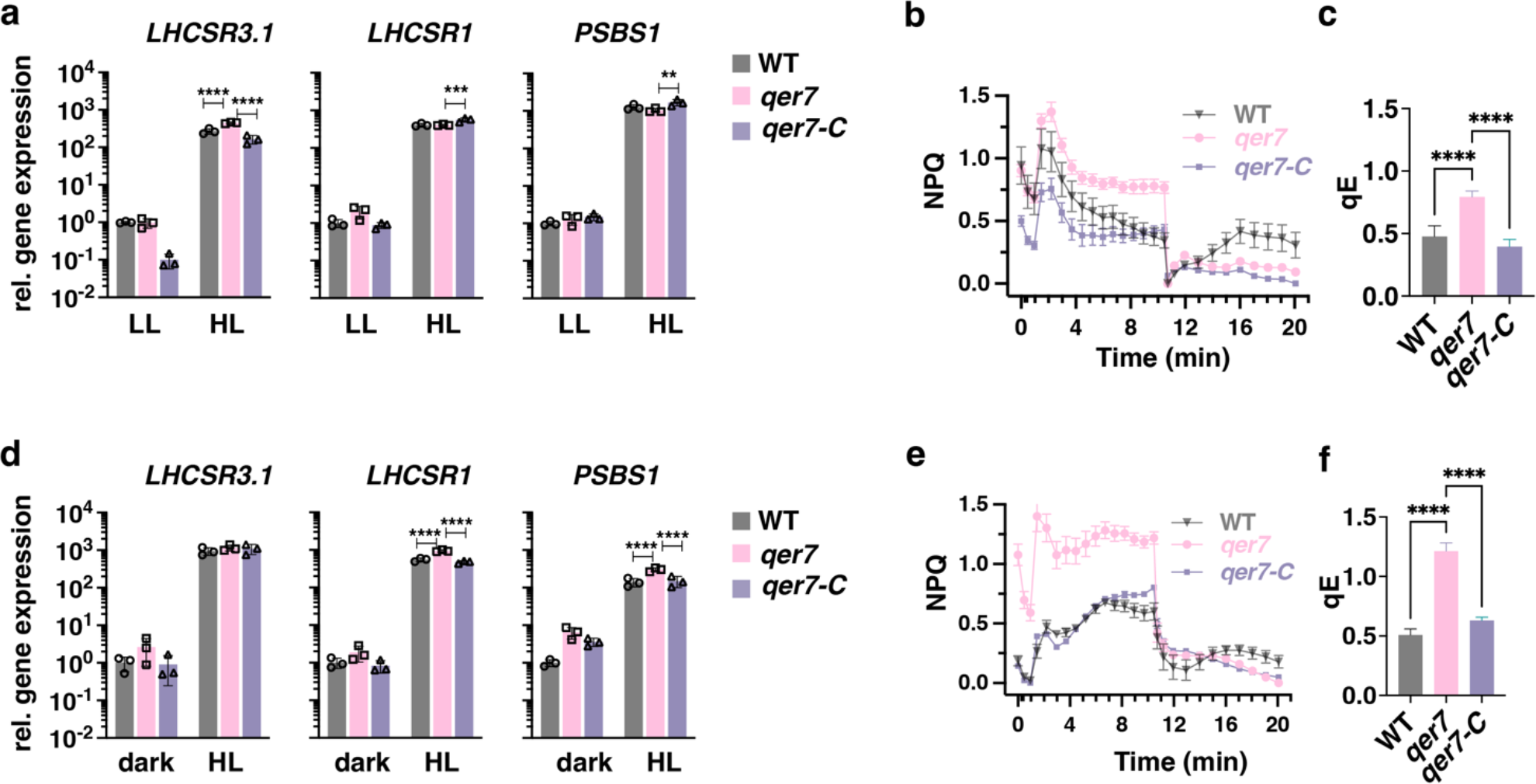
Relative expression of qE-related genes in asynchronous *qer7* cells. WT, *qer7* and *qer7-C* cells were acclimated for 16h in LL (15 µmol photons m^-2^ s^-1^; **a-c**) or darkness (**d-f**). After sampling for the LL or dark conditions, light intensity was increased to 300 µmol photons m^-2^ s^-1^ (HL) and samples were taken 1 h (RNA) or 4 h (photosynthetic measurements) after exposure to HL. **a, d**. Relative expression of qE and CCM genes at the indicated conditions normalized to WT LL (**a**) or dark (**d**) respectively (*n* = 3 biological samples, mean ± sd). **b, e.** NPQ and **c, f.** calculated qE, 4h after exposure to HL (*n* = 3 biological samples, mean ± s.d. The p-values for the comparisons are based on ANOVA Dunnett’s multiple comparisons test and are indicated in the graphs (*, P < 0.005, **, P < 0.01, ***, P < 0.001, ****, P < 0.0001). Statistical analyses for panel **a** and **d** were applied on log10- transformed values.

**Supplementary Fig. 11:**
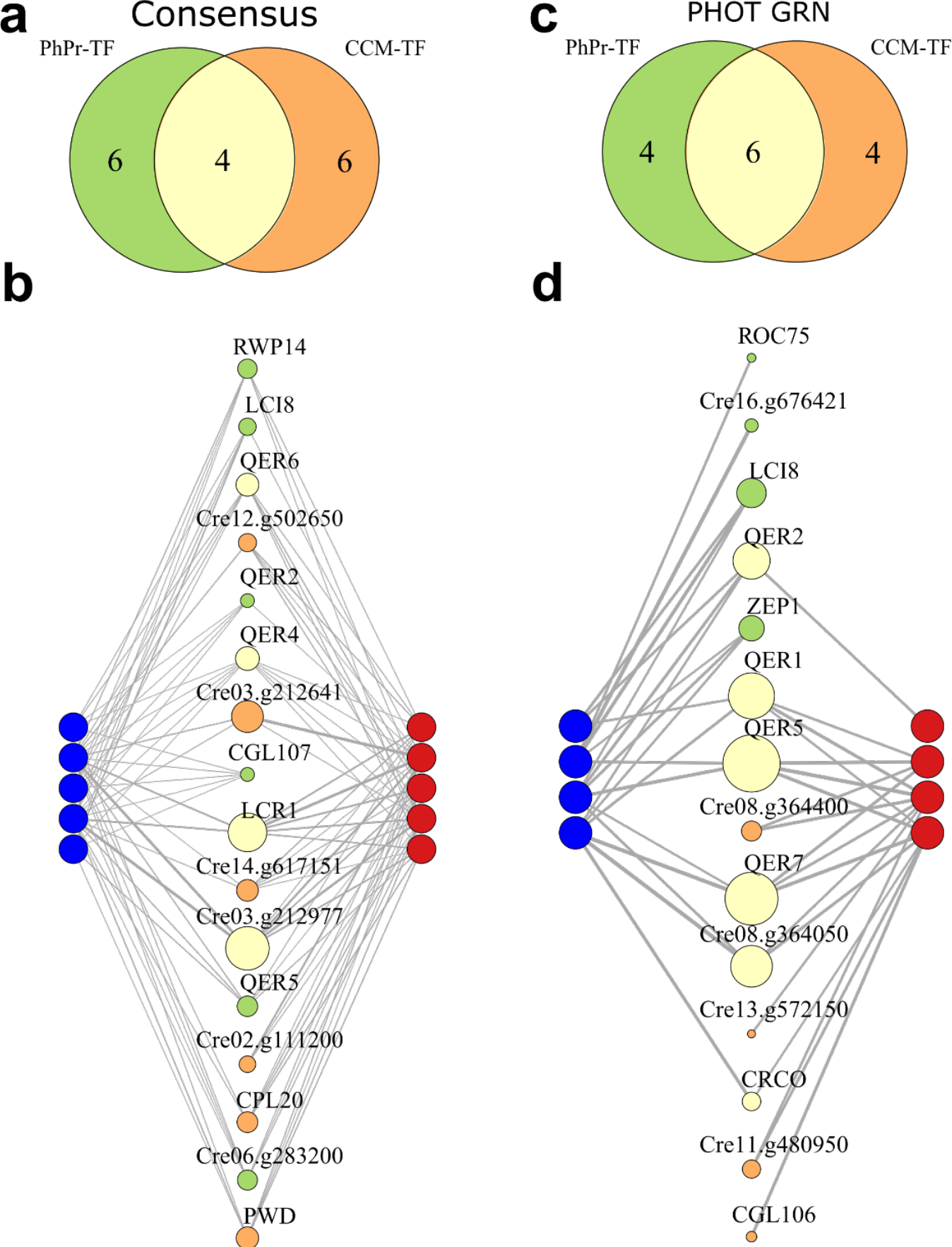
Reconstructed GRNs capture experimentally found regulators of qE and CCM. Top: Venn diagram depicting the overlap of the top 10 predicted regulators of CCM genes included in **Fig. 4a,** and qE genes based on **a.** the consensus or **c.** PHOT-specific GRN. Bottom: Network representation of the top ten TF sets of **b.** the consensus network or **d.** the phot GRN (center nodes, same color code as in panel **a**) and the target genes present in the respective network. qE-related genes (*LHCSR1*, *LHCSR3.1*, *LHCSR3.2*, *PSBS1*, *PSBS2*) are plotted in blue, and CCM genes included in **Fig. 4a** (*HLA3, CAH4, LCR1, BST1, LCI1*, *LCIA*) in red. In **b** the plotted regulatory strength corresponds to 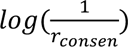, in **d**., it corresponds to the GENIE3 edge weights denoting random forest importance measure. The edge width is proportional to the strength of the specific regulatory interaction. Size of the TF nodes corresponds to the sum of all plotted target gene edge weights. See also **Supplementary Table 4,6** for QER locus IDs.

**Supplementary Fig. 12:**
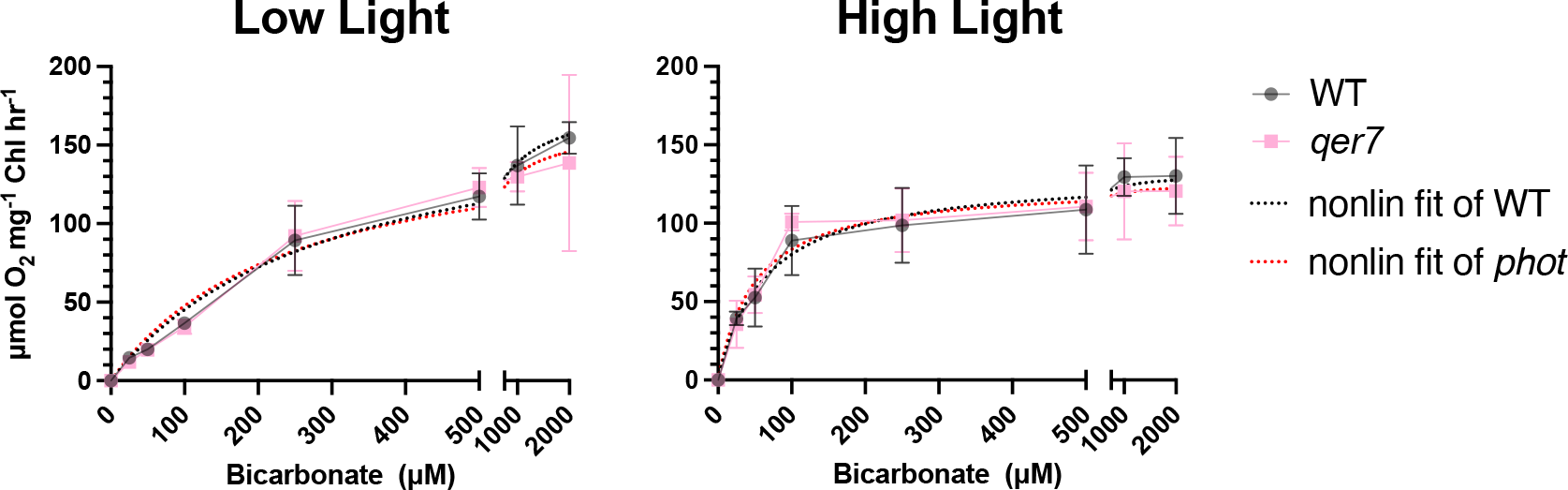
Ci affinity measurements in *qer7*. Oxygen evolution as a function of external Ci for WT and *qer7* cells grown under continuous LL (15 µmol m^-2^ s^-1^) or exposed to HL (300 µmol m^-2^ s^-1^) for 4h, in photoautotrophic conditions (Sueoka’s high salt medium; HSM), at 23 °C in Erlenmeyer flasks shaken at 125 rpm. Non-linear curve fitting to the Michaelis-Menten equation is shown in dotted lines. k1/2(Ci) and Vmax values are presented in **Supplementary Table 7**.

**Supplementary Fig. 13:**
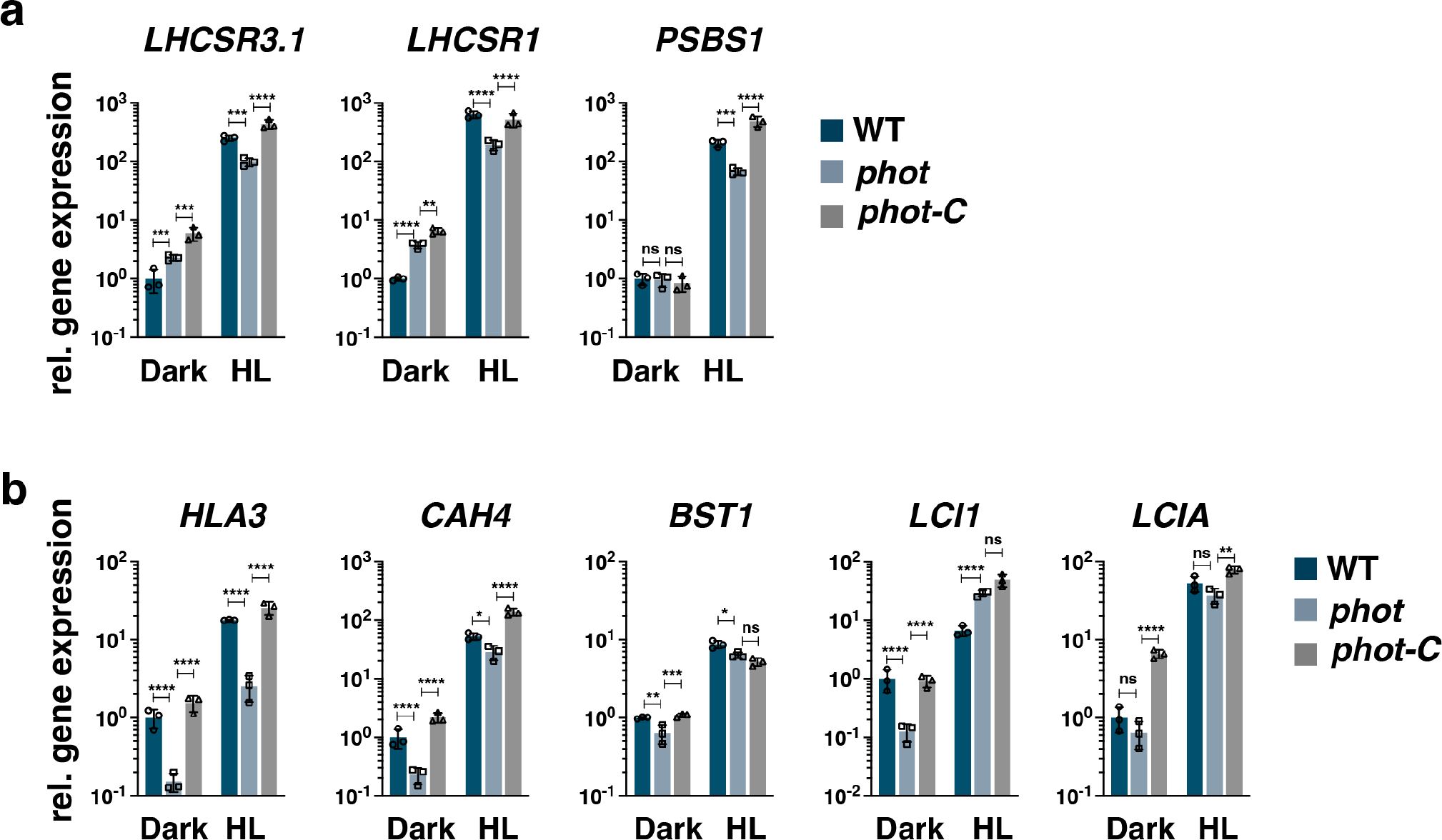
Phototropin dependent regulation of CCM gene expression. WT, *phot* and *phot-C* cells were synchronized under 12h light (15 µmol m^-2^ s^-1^)/12h dark cycles, in photoautotrophic conditions (Sueoka’s high salt medium; HSM), at 23 °C in Erlenmeyer flasks shaken at 125 rpm. After sampling at the end of the dark phase, cells were exposed to 300 µmol photons m^-2^ s^-1^ (HL) and samples were taken 1 h after HL exposure. Relative expression levels of CCM (**a**) and qE (**b**) genes at the indicated conditions normalized to WT dark (*n* = 3 biological samples, mean ± sd). The p-values for the comparisons are based on ANOVA employing Dunnett’s multiple comparisons test on log10-transformed values and are indicated in the graphs (*, P < 0.005, **, P < 0.01, ***, P < 0.001, ****, P < 0.0001).

**Supplementary Fig. 14:**
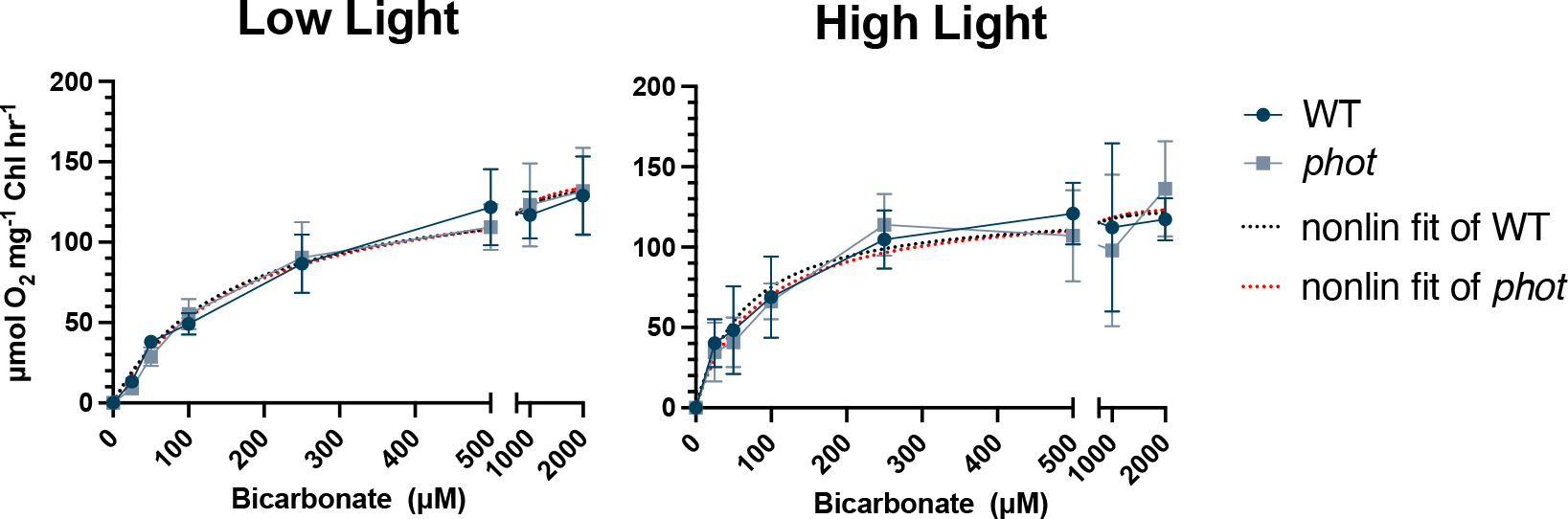
Oxygen evolution as a function of external Ci for WT and *phot* cells grown under continuous LL (15 µmol m^-2^ s^-1^) or exposed to HL (300 µmol m^-2^ s^-1^) for 4h, in photoautotrophic conditions (Sueoka’s high salt medium; HSM), at 23 °C in Erlenmeyer flasks shaken at 125 rpm. Non-linear curve fitting to the Michaelis-Menten equation is shown in dotted lines. k1/2(Ci) and Vmax values are presented in **Supplementary Table 7**.

**Supplementary Fig. 15:**
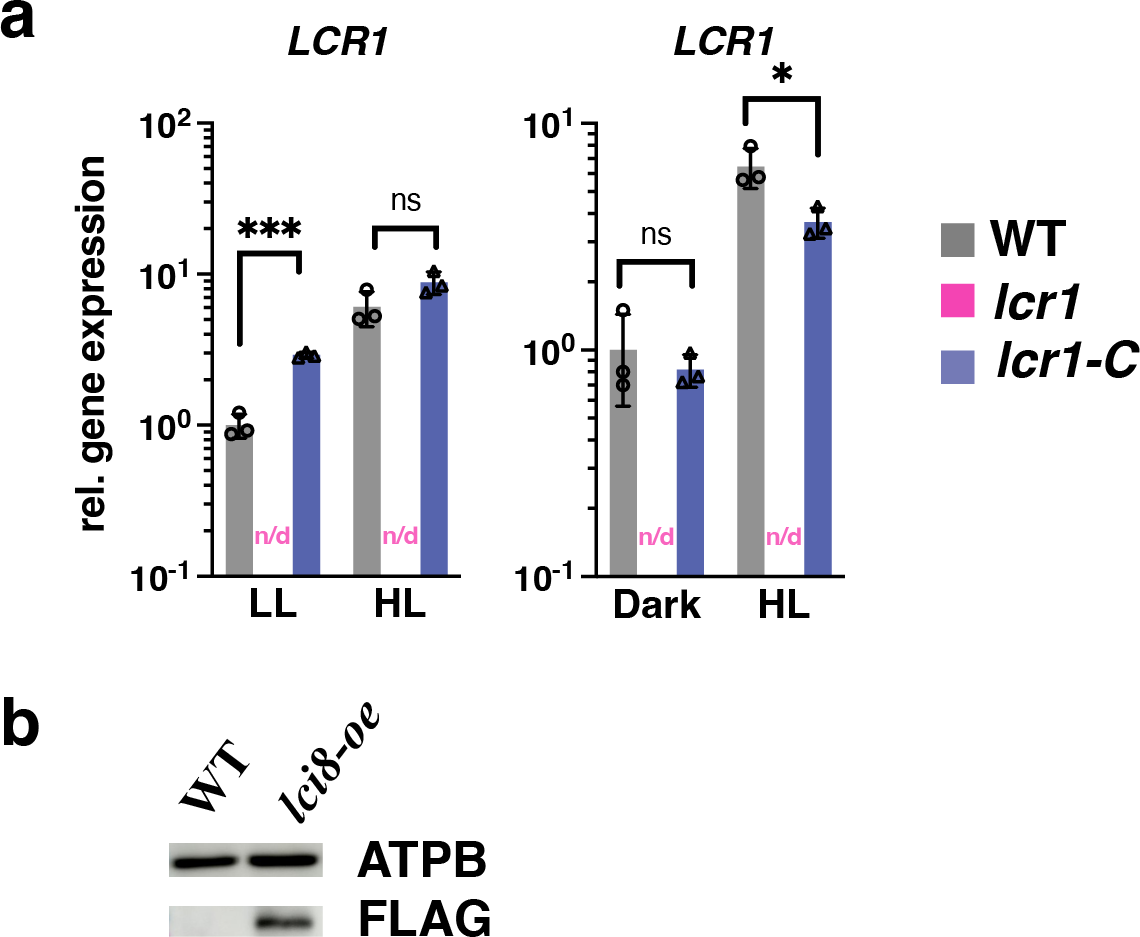
LCR1 mutation and LCI8 overexpression confirmation. **a.** Relative expression of *LCR1* in WT, *lcr1* and *lcr1-C* strains. Cells were acclimated for 16h in LL or darkness. After sampling for the LL or dark conditions, light intensity was increased to 300 µmol photons m^-2^ s^-1^ (HL), and samples for RNA purification were taken 1 h after exposure to HL (*n* = 3 biological samples, mean ± sd). *LCR1* gene expression was non- detectable (indicated as n/d in the graphs) in the *lcr1* mutant. **b. I**mmunoblot analysis of LCI8-FLAG fused protein in WT and lci8-oe cells grown in TAP, LL. The p-values for the comparisons are based on unpaired t- tests on log10- transformed values and are indicated in the graphs (*, P < 0.005, ***, P < 0.001).

## Supplementary Table Captions

Supplementary Table 1: RNAseq data sets used in this study.

Supplementary Table 2: List of Transcription factors used in this study; Family info was adapted from refs 41 and 42.

Supplementary Table 3: Edge list representation of the consensus network with mean ranks resulting from Borda count election method (refs 48 and 52) as edge attributes.

Supplementary Table 4: Top 10 predicted regulators of qE genes in the consensus GRN.

Supplementary Table 5: Edge list representation of the PHOT-specific GRN with importance score from GENIE 3 (ref. 68) as edge attributes.

Supplementary Table 6: Top 10 predicted regulators of qE genes in the PHOT-specific GRN.

Supplementary Table 7: Supplementary Table 7: k1/2(Ci) and Vmax values calculated from External Data Fig. 12 and 14.

Supplementary Table 8: List of genes putatively involved in photoprotection used for regulator prediction.

Supplementary Table 9: List of genes putatively involved in CCM used for regulator prediction.

Supplementary Table 10: Global coregulators of photoprotection and CCM based on the consensus network.

Supplementary Table 11: Global coregulators of photoprotection and CCM based on the PHOT specific network.

Supplementary Table 12: PCR primers for CLiP mutant validation and complementation in this study.

Supplementary Table 13: RT-qPCR primers for all genes in this study.

## References

1. Eberhard, S., Finazzi, G. & Wollman, F. A. The dynamics of photosynthesis. Annu. Rev. Genet. 42, 463–515 (2008).

2. Erickson, E., Wakao, S. & Niyogi, K. K. Light stress and photoprotection in Chlamydomonas reinhardtii. Plant J. 82, 449–465 (2015).

3. Moroney, J. V. & Ynalvez, R. A. Proposed carbon dioxide concentrating mechanism in Chlamydomonas reinhardtii. Eukaryotic Cell 6, 1251–1259 (2007).

4. Niyogi, K. K. PHOTOPROTECTION REVISITED: Genetic and Molecular Approaches. Annu. Rev. Plant Physiol. Plant Mol. Biol. 50, 333–359 (1999).

5. Peers, G. et al. An ancient light-harvesting protein is critical for the regulation of algal photosynthesis. Nature 462, 518–521 (2009).

6. Dinc, E. et al. LHCSR1 induces a fast and reversible pH-dependent fluorescence quenching in LHCII in Chlamydomonas reinhardtii cells. Proc. Natl. Acad. Sci. U.S.A. 113, 7673–7678 (2016).

7. Ruiz-Sola, M. et al. Photoprotection is regulated by light-independent CO2 availability. bioRxiv (2021). Doi:10.1101/2021.10.23.465040

8. Li, X. P. et al. A pigment-binding protein essential for regulation of photosynthetic light harvesting. Nature 403, 391–395 (2000).

9. Correa-Galvis, V. et al. Photosystem II Subunit PsbS Is Involved in the Induction of LHCSR Protein- dependent Energy Dissipation in Chlamydomonas reinhardtii. J. Biol. Chem. 291, 17478–17487 (2016).

10. Tibiletti, T., Auroy, P., Peltier, G. & Caffarri, S. Chlamydomonas reinhardtii PsbS Protein Is Functional and Accumulates Rapidly and Transiently under High Light. Plant Physiol. 171, 2717–2730 (2016).

11. Allorent, G. et al. UV-B photoreceptor-mediated protection of the photosynthetic machinery in Chlamydomonas reinhardtii. Proc. Natl. Acad. Sci. U.S.A. 113, 14864–14869 (2016).

12. Redekop, P. et al. PsbS contributes to photoprotection in Chlamydomonas reinhardtii independently of energy dissipation. Biochimica et Biophysica Acta (BBA) – Bioenergetics 1861, 148183 (2020).

13. Hanawa, Y., Watanabe, M., Karatsu, Y., Fukuzawa, H. & Shiraiwa, Y. Induction of a high-CO2- inducible, periplasmic protein, H43, and its application as a high-CO2-responsive marker for study of the high-CO2-sensing mechanism in Chlamydomonas reinhardtii. Plant Cell Physiol. 48, 299–309 (2007).

14. Moroney, J. V. et al. Isolation and Characterization of a Mutant of Chlamydomonas reinhardtii Deficient in the CO2 Concentrating Mechanism. Plant Physiol. 89, 897–903 (1989).

15. Fukuzawa, H. et al. Ccm1, a regulatory gene controlling the induction of a carbon-concentrating mechanism in Chlamydomonas reinhardtii by sensing CO2 availability. Proc. Natl. Acad. Sci. U.S.A. 98, 5347–5352 (2001).

16. Xiang, Y., Zhang, J. & Weeks, D. P. The Cia5 gene controls formation of the carbon concentrating mechanism in Chlamydomonas reinhardtii. Proc. Natl. Acad. Sci. U.S.A. 98, 5341–5346 (2001).

17. Yoshioka, S. et al. The novel Myb transcription factor LCR1 regulates the CO2-responsive gene Cah1, encoding a periplasmic carbonic anhydrase in Chlamydomonas reinhardtii. Plant Cell 16, 1466–1477 (2004).

18. Petroutsos, D. et al. A blue-light photoreceptor mediates the feedback regulation of photosynthesis. Nature 537, 563–566 (2016).

19. Petroutsos, D. et al. The chloroplast calcium sensor CAS is required for photoacclimation in *Chlamydomonas reinhardtii*. Plant Cell 23, 2950–2963 (2011).

20. Maruyama, S., Tokutsu, R. & Minagawa, J. Transcriptional regulation of the stress-responsive light harvesting complex genes in *Chlamydomonas reinhardtii*. Plant Cell Physiol. 55, 1304–1310 (2014).

21. Miura, K. et al. Expression profiling-based identification of CO2-responsive genes regulated by CCM1 controlling a carbon-concentrating mechanism in Chlamydomonas reinhardtii. Plant Physiol. 135, 1595–1607 (2004).

22. Fang, W. et al. Transcriptome-wide changes in Chlamydomonas reinhardtii gene expression regulated by carbon dioxide and the CO2-concentrating mechanism regulator CIA5/CCM1. Plant Cell 24, 1876–1893 (2012).

23. Brueggeman, A. J. et al. Activation of the carbon concentrating mechanism by CO2 deprivation coincides with massive transcriptional restructuring in Chlamydomonas reinhardtii. Plant Cell 24, 1860–1875 (2012).

24. Aihara, Y., Fujimura-Kamada, K., Yamasaki, T. & Minagawa, J. Algal photoprotection is regulated by the E3 ligase CUL4-DDB1DET1. Nature Plants 5, 34–40 (2019).

25. Redekop, P. et al. Transcriptional regulation of photoprotection in dark-to-light transition- more than just a matter of excess light energy. Sci Adv (2022).

26. Gabilly, S. T. et al. Regulation of photoprotection gene expression in Chlamydomonas by a putative E3 ubiquitin ligase complex and a homolog of CONSTANS. Proc. Natl. Acad. Sci. U.S.A. 116, 17556– 17562 (2019).

27. Tokutsu, R., Fujimura-Kamada, K., Matsuo, T., Yamasaki, T. & Minagawa, J. The CONSTANS flowering complex controls the protective response of photosynthesis in the green alga Chlamydomonas. Nat Comms 10, 655–10 (2019).

28. Kamrani, Y. Y., Matsuo, T., Mittag, M. & Minagawa, J. ROC75 is an Attenuator for the Circadian Clock that Controls LHCSR3 Expression. Plant Cell Physiol. 59, 2602–2607 (2018).

29. Omranian, N., Eloundou-Mbebi, J. M. O., Mueller-Roeber, B. & Nikoloski, Z. Gene regulatory network inference using fused LASSO on multiple data sets. Sci Rep 6, 1–14 (2016).

30. Marbach, D. et al. Wisdom of crowds for robust gene network inference. Nat. Methods 9, 796–804 (2012).

31. Fang, L., et al. GRNdb: Decoding the gene regulatory networks in diverse human and mouse conditions. Nucl. Acids Res. 49, D97–D103 (2021).

32. Omranian, N. & Nikoloski, Z. in Methods in Molecular Biology 1629, 283–295 (2017).

33. Huynh-Thu, V. A. & Sanguinetti, G. in Methods in Molecular Biology 1883, 1–23 (2019).

34. Schäfer, J. & Strimmer, K. A shrinkage approach to large-scale covariance matrix estimation and implications for functional genomics. Stat Appl Genet Mol Biol 4, Article32 (2005).

35. Zou, H. & Hastie, T. Regularization and variable selection via the elastic net. Journal of the Royal Statistical Society: Series B (Statistical Methodology) 67, 301–320 (2005).

36. Gargouri, M. et al. Identification of regulatory network hubs that control lipid metabolism in Chlamydomonas reinhardtii. J. Exp. Bot. 66, 4551–4566 (2015).

37. Lämmermann, N., et al. Ubiquitin ligase component LRS1 and transcription factor CrHy5 act as a light switch for photoprotection in *Chlamydomonas*. bioRxiv 2020.02.10.942334 (2020).

38. Zones, J. M., Blaby, I. K., Merchant, S. S. & Umen, J. G. High-Resolution Profiling of a Synchronized Diurnal Transcriptome from Chlamydomonas reinhardtii Reveals Continuous Cell and Metabolic Differentiation. Plant Cell 27, 2743–2769 (2015).

39. Strenkert, D. et al. Multiomics resolution of molecular events during a day in the life of Chlamydomonas. Proc. Natl. Acad. Sci. U.S.A. 116, 2374–2383 (2019).

40. Fett, J. P. & Coleman, J. R. Regulation of Periplasmic Carbonic Anhydrase Expression in Chlamydomonas reinhardtii by Acetate and pH. Plant Physiol. 106, 103–108 (1994).

41. Pérez-Rodríguez, P. et al. PlnTFDB: Updated content and new features of the plant transcription factor database. Nucl. Acids Res. 38, (2009).

42. Jin, J. et al. PlantTFDB 4.0: Toward a central hub for transcription factors and regulatory interactions in plants. Nucl. Acids Res. 45, D1040–D1045 (2017).

43. Fischer, B. B. et al. SINGLET OXYGEN RESISTANT 1 links reactive electrophile signaling to singlet oxygen acclimation in Chlamydomonas reinhardtii. Proc. Natl. Acad. Sci. U.S.A. 109, E1302–11 (2012).

44. Margolin, A. A. et al. ARACNE: An algorithm for the reconstruction of gene regulatory networks in a mammalian cellular context. BMC Bioinformatics 7, 1–15 (2006).

45. Barzel, B. & Barabási, A.-L. Network link prediction by global silencing of indirect correlations. Nature Biotechnology 31, 720–725 (2013).

46. Omranian, N., Eloundou-Mbebi, J. M. O., Mueller-Roeber, B. & Nikoloski, Z. Gene regulatory network inference using fused LASSO on multiple data sets. Sci Rep 6, 20533 (2016).

47. Cerulo, L., Elkan, C. & Ceccarelli, M. Learning gene regulatory networks from only positive and unlabeled data. BMC Bioinformatics 11, 228–16 (2010).

48. de Borda, J.-C. Mémoire sur les élections au scrutin. Histoire de l’Academie Royale des Sciences (1781).

49. Li, X. et al. A genome-wide algal mutant library and functional screen identifies genes required for eukaryotic photosynthesis. Nat. Genet. 51, 627–635 (2019).

50. Redekop, P. et al. Transcriptional regulation of photoprotection in dark-to-light transition-More than just a matter of excess light energy. Sci Adv 8, (2022).

51. UniProt Consortium. UniProt: the universal protein knowledgebase in 2021. Nucleic Acids Res. 49, D480–D489 (2021).

52. Marbach, D. et al. Wisdom of crowds for robust gene network inference. Nat. Methods 9, 796–804 (2012).

53. Mi, H. et al. PANTHER version 16: A revised family classification, tree-based classification tool, enhancer regions and extensive API. Nucl. Acids Res. 49, D394–D403 (2021).

54. Thiriet-Rupert, S. et al. Transcription factors in microalgae: genome-wide prediction and comparative analysis. BMC Genomics 17, 282 (2016).

55. Yoshioka, S. et al. The novel Myb transcription factor LCR1 regulates the CO2-responsive gene Cah1, encoding a periplasmic carbonic anhydrase in Chlamydomonas reinhardtii. Plant Cell 16, 1466–1477 (2004).

56. Borodin, V. B. Effect of red and blue light on acclimation of Chlamydomonas reinhardtii to CO2- limiting conditions. Russian Journal of Plant Physiolofy 55, 441–448 (2008).

57. Santhanagopalan, I., Wong, R., Mathur, T. & Griffiths, H. Orchestral manoeuvres in the light: crosstalk needed for regulation of the Chlamydomonas carbon concentration mechanism. J. Exp. Bot. 72, 4604–4624 (2021).

58. Chaux, F., Peltier, G. & Johnson, X. A security network in PSI photoprotection: regulation of photosynthetic control, NPQ and O2 photoreduction by cyclic electron flow. Front Plant Sci 6, 875 (2015).

59. Choi, B. Y. et al. The Chlamydomonas bZIP transcription factor BLZ8 confers oxidative stress tolerance by inducing the carbon-concentrating mechanism. Plant Cell 34, 910–926 (2022).

60. Scholz, M. et al. Light-dependent N-terminal phosphorylation of LHCSR3 and LHCB4 are interlinked in Chlamydomonas reinhardtii. Plant J. 99, 877–894 (2019).

61. Wang, L. et al. Chloroplast-mediated regulation of CO2-concentrating mechanism by Ca2+-binding protein CAS in the green alga Chlamydomonas reinhardtii. Proc. Natl. Acad. Sci. U.S.A. 113, 12586– 12591 (2016).

62. Hiyama, A. et al. Blue light and CO2 signals converge to regulate light-induced stomatal opening. Nat Comms 8, 1284 (2017).

63. Bushnell, B. BBMap: A Fast, Accurate, Splice-Aware Aligner. *Lawrence Berkeley National Laboratory*. LBNL Report LBNL-E (2014).

64. Law, C. W., Chen, Y., Shi, W. & Smyth, G. K. Voom: Precision weights unlock linear model analysis tools for RNA-seq read counts. Genome Biology 15, 1–17 (2014).

65. Anders, S., Pyl, P. T. & Huber, W. HTSeq-A Python framework to work with high-throughput sequencing data. Bioinformatics 31, 166–169 (2015).

66. Robinson, M. D. & Oshlack, A. A scaling normalization method for differential expression analysis of RNA-seq data. Genome Biology 11, 1–9 (2010).

67. Robinson, M. D., McCarthy, D. J. & Smyth, G. K. edgeR: A Bioconductor package for differential expression analysis of digital gene expression data. Bioinformatics 26, 139–140 (2009).

68. Huynh-Thu, V. A., Irrthum, A., Wehenkel, L. & Geurts, P. Inferring regulatory networks from expression data using tree-based methods. PloS ONE 5, e12776 (2010).

69. Meyer, P. E., Lafitte, F. & Bontempi, G. Minet: A r/25einhardtii25 package for inferring large transcriptional networks using mutual information. BMC Bioinformatics 9, 1–10 (2008).

70. Olsen, C., Meyer, P. E. & Bontempi, G. On the impact of entropy estimation on transcriptional regulatory network inference based on mutual information. Eurasip Journal on Bioinformatics and Systems Biology 2009, (2009).

71. Faith, J. J. et al. Large-scale mapping and validation of Escherichia coli transcriptional regulation from a compendium of expression profiles. PloS Biol. 5, 0054–0066 (2007).

72. Feizi, S., Marbach, D., Médard, M. & Kellis, M. Network deconvolution as a general method to distinguish direct dependencies in networks. Nature Biotechnology 31, 726–733 (2013).

73. Ritchie, M. E., et al. Limma powers differential expression analyses for RNA-sequencing and microarray studies. Nucl. Acids Res. 43, e47 (2015).

74. Love, M. I., Huber, W. & Anders, S. Moderated estimation of fold change and dispersion for RNA-seq data with DESeq2. Genome Biology 15, 1–21 (2014).

75. North, B. V., Curtis, D. & Sham, P. C. A note on the calculation of empirical P values from Monte Carlo procedures [1]. American Journal of Human Genetics 71, 439–441 (2002).

76. Gorman, D. S. & Levine, R. P. Cytochrome f and plastocyanin: their sequence in the photosynthetic electron transport chain of *Chlamydomonas 25einhardtii*. Proc. Natl. Acad. Sci. U.S.A. 54, 1665–1669 (1965).

77. Sueoka, N. Mitotic replication of deoxyribonucleic acid in *Chlamydomonas 25einhardtii*. Proc. Natl. Acad. Sci. U.S.A. 46, 83–91 (1960).

78. Greiner, A. et al. Targeting of Photoreceptor Genes in Chlamydomonas reinhardtii via Zinc-Finger Nucleases and CRISPR/Cas9. Plant Cell 29, 2498–2518 (2017).

79. Kaye, Y. et al. The mitochondrial alternative oxidase from Chlamydomonas reinhardtii enables survival in high light. J. Biol. Chem. 294, 1380–1395 (2019).

80. Gibson, D. G. et al. Enzymatic assembly of DNA molecules up to several hundred kilobases. Nat. Methods 6, 343–345 (2009).

81. Mackinder, L. C. M. et al. A repeat protein links Rubisco to form the eukaryotic carbon-concentrating organelle. Proc. Natl. Acad. Sci. U.S.A. 113, 5958–5963 (2016).

82. Schloss, J. A. A Chlamydomonas gene encodes a G protein beta subunit-like polypeptide. Molec. Gen. Genet. 221, 443–452 (1990).

83. Schlesselman, J. J. Data Transformation in Two-Way Analysis of Variance. Journal of the American Statistical Association (2012). Doi:10.1080/01621459.1973.10482435

84. Genty, B., Briantais, J.-M. & Baker, N. R. The relationship between the quantum yield of photosynthetic electron transport and quenching of chlorophyll fluorescence. Biochimica et Biophysica Acta (BBA) 990, 87–92 (1989).

85. Mukherjee, A. et al. Thylakoid localized bestrophin-like proteins are essential for the CO2 concentrating mechanism of Chlamydomonas reinhardtii. Proc. Natl. Acad. Sci. U.S.A. 116, 16915–16920 (2019).

## Supplementary list of references

1. Li, X. et al. A genome-wide algal mutant library and functional screen identifies genes required for eukaryotic photosynthesis. Nat. Genet. 51, 627–635 (2019).

2. Marbach, D. et al. Wisdom of crowds for robust gene network inference. Nat. Methods 9, 796–804 (2012).

